# Bone marrow stromal cell transplantation ameliorates cytopenia caused by depletion FAP-α expressing cells

**DOI:** 10.1101/2020.11.23.393959

**Authors:** Eva Camarillo-Retamosa, Luke Watson, Paul Loftus, Senthil Alagesan, Yvonne O’Donoghue, Laura M. Deedigan, Lisa O’Flynn, Laura R. Barkley, Stephen J.Elliman

**Affiliations:** Orbsen Therapeutics Ltd., Galway, Ireland; School of Medicine, College of Medicine, Nursing and Health Sciences, National University of Ireland, Galway, Ireland; Lambe Institute for Translational Research, National University of Ireland Galway, Ireland; Avectas Ltd, Maynooth University, Co. Kildare, Ireland

**Keywords:** Fibroblast activation protein-α (FAP), transgenic mouse model (FAPDM2), Diphtheria Toxin (DTX), Peripheral Blood (PB), Bone marrow (BM), Mesenchymal stromal cells (MSC), Hematopoietic stem cells (HSC)

## Abstract

The marrow microenvironment is a complex and heterogeneous mixture of hematopoietic and stromal progenitors necessary for haematopoiesis. Whilst the hematopoietic progenitors are well described, the stromal cellular composition is not fully elucidated due to the low cells numbers, localisation-distribution-accessibility, and the lack of specific biomarkers. Cellular taxonomy studies have recently identified new populations of stromal subsets with distinct gene signature and regulatory properties of hematopoietic regeneration. Fibroblast activation protein-α (FAP), a stromal cell type first identified in cancer is also rarely found in normal tissues but might play an essential role in tissue homeostasis. Using FAP^DM2^ transgenic mouse in which FAP-expressing cells can be ablated with Diphtheria Toxin (DTX) FAP^+^ cells were depleted in healthy mice. Whilst FAP^+^ cells constituted 5% of all marrow cells; its ablation caused a rapid loss of PDGFR-α, Leptin-R, gp38 and SDC2 stromal cells populations, endothelial cells and vascular disruption. These resulted in anaemia, thrombocytopenia and neutropenia in peripheral blood (PB) and extreme hypo-cellularity in marrow with abnormalities within the hematopoietic progenitors. In an effort to reverse the phenotypes caused by FAP^+^ cell loss, a single intravenous injection of syngeneic bone marrow-derived stromal cells was administered. In a short-term evaluation, anaemia, thrombocytopenia and neutropenia ameliorated in PB and the numbers of marrow hematopoietic progenitors increased. Our data suggest FAP-expressing cells are a non-redundant component of the marrow microenvironment, necessary for marrow homeostasis and haematopoiesis. These data also provided evidence that stromal cell ablation can be rescued by stromal cell therapy.

**Significance Statement:** FAP-expressing cells depletion led to collateral damage in PB and marrow, including haematological defects that can be ameliorated by adoptive transfer of low-dose, *ex-vivo* expanded FAP-expressing marrow stromal cells. We suggest that stromal cell loss is a feature of severe immune-mediated inflammatory diseases – such as Graft versus Host Disease and sepsis - and that FAP^DM2^ model represents a novel tool to explore the native function of the recently identified stromal cell sub-populations.

## Introduction

The paucity and heterogeneity of the stromal cells, together with their eventual limited access and technical challenges on cellular identity, resulting in a poorly defined non-parenchymal element of every tissue with critical roles in organ development, homeostasis, and tissue repair. In consequence, the understanding of the cell-based therapies efficacy is limited. Recent studies of single-cell RNA sequencing have identified 17 clusters of 6 stromal cell subpopulations in mouse marrow – each with a distinct signature^1^.

Within the distinctive markers, FAP^+^ stromal cells were first demonstrated in human adenocarcinomas and subsequently were found in non-cancerous inflammatory lesions ^2, 3,^ Recently, a genetically modified mouse model in which FAP^+^ cells are expressing the primate diphtheria toxin receptor (DTXR) was generated in which the conditional depletion of these cells from an established, immunogenic, a transplanted tumour caused its growth arrest ^4^. Notably, in the same DTXR model, experimental ablation of these FAP^+^ cells causes loss of muscle mass and a reduction of lymphopoiesis and erythropoiesis, revealing their essential functions in maintaining average muscle mass and haematopoiesis, respectively^5, 6^. FAP is a membrane-bound protease found in healthy bone marrow, umbilical cord and adipose tissues^7^. Osteolineage cells expressing FAP overlaps with PDGFR-α^8^ and Leptin-R^9, 10^ supporting bone turnover in the bone formation and endothelial vascular homeostasis in haematopoiesis within the BM^1, 5, 8, 11^. FAP^+^ cells participate in haematopoiesis producing stromal cell factors cxcl12 and Kit-L^5^, but its signature in homeostasis and physiological roles are not well defined due to their scarcity and heterogeneity in normal tissues^1, 12-15^.

More recently, FAP^+^ stromal cells have shown (1) to behave as immune effector cells associated with tissue damage in models of immune-mediated arthritis^16^ and (2) that embryonic FAP^+^ cells have ontogenic origins in lymph node stromal cells^17^. Using FAP^DM2^ model, we analysed the impact of FAP^+^ cell ablation on the PB and BM in normal healthy mice. We sought to investigate if a single intravenous transference of syngeneic marrow stromal cells could repair or ameliorate the haematological and stromal damage elicited by FAP^+^ cell ablation. We present the FAP^DM2^ model as an *in vivo* platform to investigate the function of the novel populations of stromal cells identified.

## Methods

Detailed methods are shown in the supplemental information.

## Results

### FAP^+^ cell ablation eliminated the stromal and endothelial compartments in BM after 72 hours

We administered two doses of DTX in FAP transgenic (FAP) and littermate (LM) mice to ablate *fap*^+^ cells and evaluate the BM cellularity at 72hrs (D3) (Figure 1A, supplemental table 1 and figure1). Flow cytometry analysis indicated that DTX-administration deleted the 82% of the stromal compartment (CD45^−^CD31^−^) (Figure 1B) and the 75% of endothelial cells (CD45^−^ CD31^+^) in FAP (Figure 1E, F, supplemental table 3 and figure 7) when compared to LM 22 days after DTX-administration (D22) (Figure 1C). Data from our studies in digested femurs demonstrated that complexity of the BM cellularity is comparable between LM and FAP (Figure 1D) including stromal (10%), endothelial (>3%), haematological (>50%) or the double-positive (>30%) cellular subsets (Figure 1E, supplemental table 3 and figure 2). The stromal cell population expressing FAP (<2×10^4^) represents the 0.05% of the whole marrow confirming its scarcity. However, it represents the largest cluster when compared to those expressing gp38, SDC2, PDGFR-α or Leptin-R, which were all also ablated (Figure 1F, supplemental table 4, figures 2 and 8).

**Figure 1.**
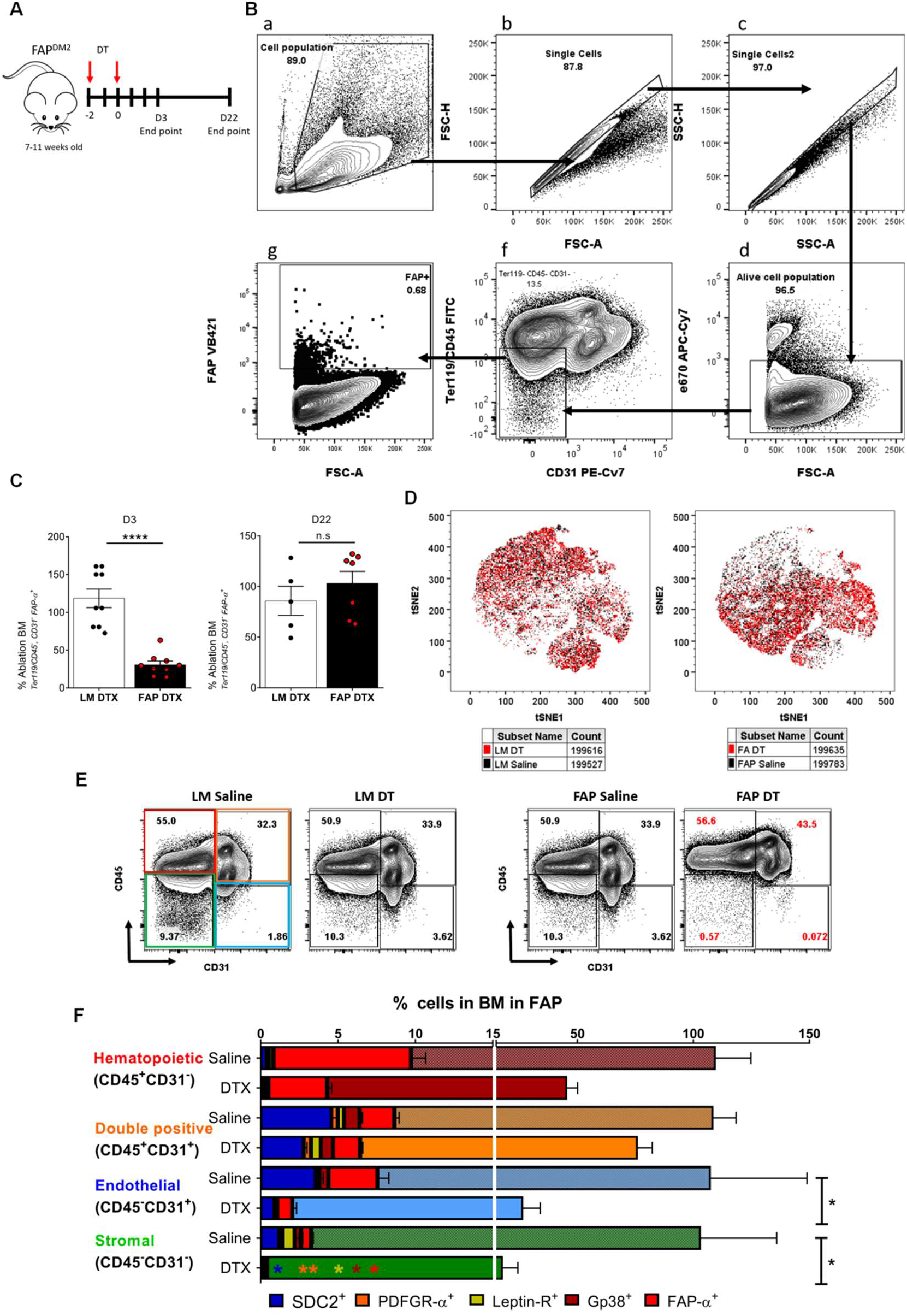
DTX-mediated FAP-cell ablation caused rapid loss of multiple stromal cells populations, endothelial and haematopoietic cell types. **(A)** Schematic workflow of the DTX injection regime and endpoint analysis. **(B)** Representative flow cytometry gating strategy to examine the stromal cell population (Ter119^−^, CD45^−^, CD31^−^) (f) expressing FAP (g) in a bone marrow sample. **(C)** Frequency of stromal cell population expressing FAP (Ter119-CD45-CD31-, FAP+) after DTX administration. D3 LM-DTX and FAP-DTX (n=9 per group), D22 LM-DTX (n=5) FAP-DTX (n=8) α=0.05, Mean ± SEM;***,P<0.001. **(D)** t-SNE visualization of 200,000 nucleated cells from a representative bone marrow sample from LM overlaid with FAP sample after DTX administration. **(E)** Representative flow cytometry dot plots with colour-coded squares for bone marrow cell clusters based on the expression of CD45 and CD31 in LM and FAP after DTX administration. **(F)** Frequency of mouse bone marrow cells within each colour-coded cluster after DTX administration in FAP. Multi-grouped bar graph referred to the cell number obtained with the saline control. n=4 per group, mean ± SEM; α=0.05, *,P<0.05.

**Figure 2.**
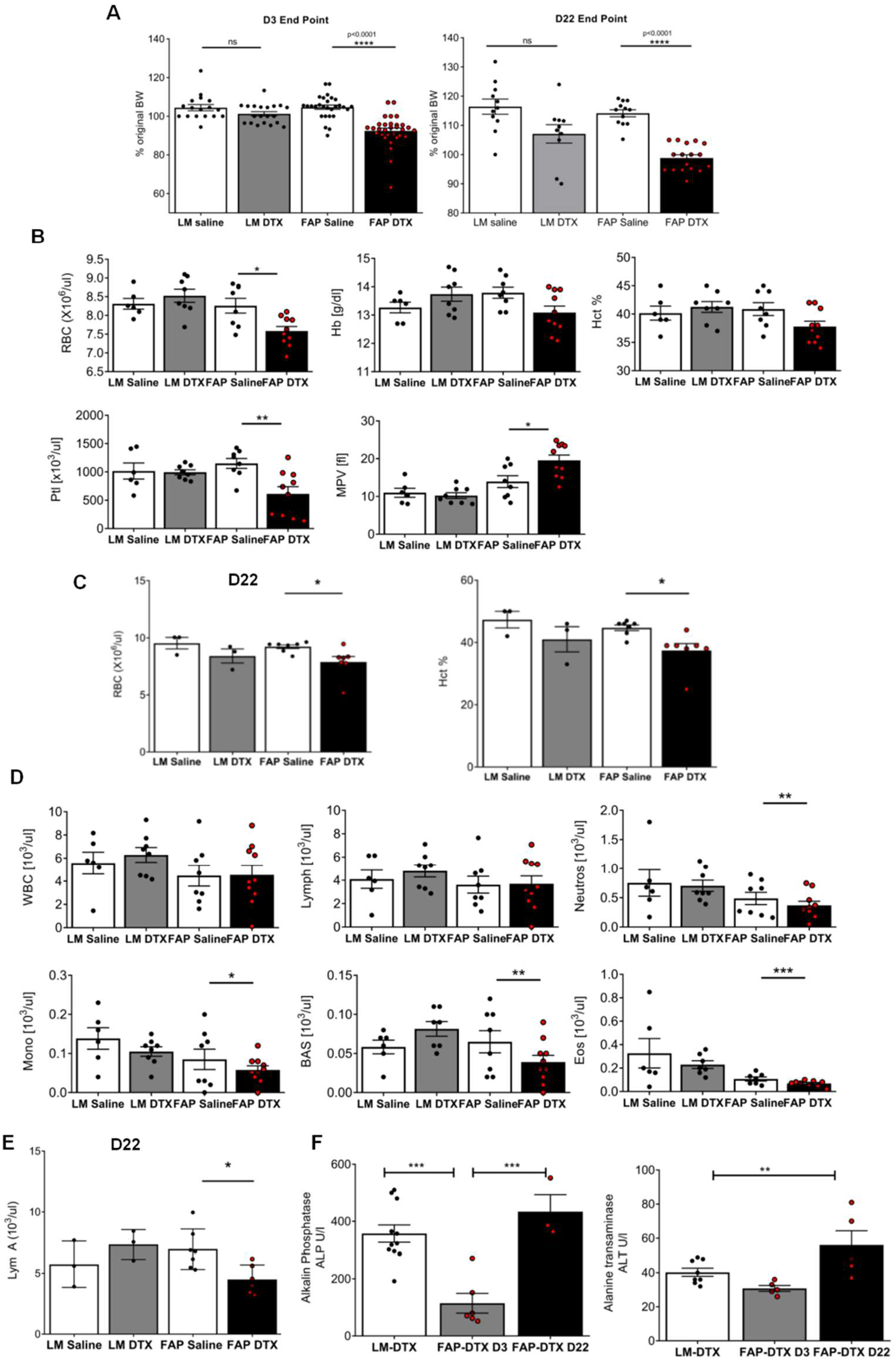
DTX-mediated FAP-cell ablation caused bodyweight loss and cytopenia in peripheral blood. **(A)** Percentage of the original body weight (BW) at D33 and D22 after DTX administration. LM-saline (n=17), LM-DTX (n=20), FAP-Saline (n=18), FAP-DTX (n=31) for D3 and LM- saline (n=11), LM-DTX (n=10), FAP-Saline (n=11), FAP-DTX (n=17) for D22. Mean ± SEM, α=0.05 ****,P<0.0001. **(B)** Red blood parameters from peripheral blood at D33 after DTX administration. LM-saline (n=6), LM-DTX (n=8), FAP-Saline (n=11), FAP-DTX (n=10) Mean ± SEM; α=0.05 *,P<0.05; **,P<0.01). **(C)** The amount of red blood cells at D22 after DTX administration. LM-saline and DTX (n=3), FAP-Saline and DTX (n=7), α=0.05, Mean ± SEM; *,P<0.05; **,P<0.01. **(D)** White blood parameters from peripheral blood at D3 after DTX administration. LM-saline (n=6), LM-DTX (n=8), FAP-Saline (n=11), FAP-DTX (n=10), α=0.05, Mean ± SEM; *,P<0.05; **,P<0.01. **(E)** Absolute number of lymphocytes in peripheral blood at D22 after DTX administration. LM-saline and DTX (n=3), FAP-Saline and DTX (n=7), α=0.05, Mean ± SEM; *,P<0.05. **(F)** Biochemistry parameters in serum obtained at D3 and D22 after DTX administration. LM-DTX (n=8-11), FAP-DTX D3 (n=3-6) and FAP-DTX D22 (n=1-5), α=0.05, Mean ± SEM. **,P<0.01, ***,P<0.001

A t-distributed stochastic neighbour embedding (t-SNE) analysis of 200,000 events from a representative sample of FAP^DM2^ identified the cellular diversity and distribution (Supplemental Figure 2) based on CD45 and CD31 expression: stromal (CD45^−^CD31^−^), endothelial (CD45^−^CD31^+^), haematological (CD45^+^CD31^−^), double-positive (CD45^+^CD31^+^) (Supplemental Figure 2II). We also identified the widely distributed FAP, SDC2, PDGFR-α, Leptin-R and gp38 stromal cell markers in the four-compartments (Supplemental Figure 2 GIII). A recent analysis on scRNA-seq^1^ allowed us to confirm the ubiquitous presence of *fap* at a molecular level which is highly expressed in osteogenic progenitor cluster (OLC-1), chondrogenic progenitor (2&4) and fibroblastic clusters (Supplemental figure 3). Moreover, *fap* co-localised with *sdc2, pdgfrα* and, to a lesser degree, with *lepr* in OLC-1 and chondro-progenitors clusters (2&4). In contrast, few fibroblasts expressed *fap* in homeostasis, which overlaps with *pdpn (gp38)* (Supplemental figure 4). Interestingly, *cxcl12* and *csf1* factors were both highly expressed in a small proportion of OLC-1 and scarce in fibroblast cluster 9, and *kitl* expression was much lower. Notably, the expression of *il7* factor, recently linked to bone homeostasis^18^, was restricted to a small proportion of OLC-1. This comprehensive analysis was required as the ablative therapy has profound implications in the whole organism^16^.

### FAP^+^ cell ablation causes anaemia, thrombocytopenia and granulopenia

FAP^+^ cell ablation caused a 10% loss of body weight in FAP that did not recover by D22 (Figure 2A). To evaluate the overall health, we analysed the blood counts. The cellular size, maturity (RDW and MPV), and the amount of haemoglobin carried by erythrocytes (MHC and MHCH) confirming an anaemia phenotype^5^ (Figure 2B). The blood amount (%Htc) and the total number of erythrocytes (RBC) were both reduced by 5% and 7% respectively resulting in a 2.84% decrease of Hb which did not resolve at D22 (Figure 2C). Platelets were reduced in number (45.13%) although the remaining ones were 50% bigger in size (MPV) (Figure 2B). The immune cells reduced by 14% (WBC) (Figure 2D) including neutrophils, monocytes, basophils and eosinophils whilst lymphocytes dropped 24.9% at D22 (Figure 2E). Liver and renal parameters responsible for erythrocytes turnover tested in serum confirming DTX toxicity to *fap-*DTXR transgene (Data not shown). Liver dysfunction enzymes ALP (alkaline phosphatase) - responsible for removing phosphate groups (de-phosphorylation) - and ALT (alanine aminotransferase) – a trigger for tissue damage or disease - were both reduced by 67.85% and 23.47% respectively at D3 whilst the levels increased by 21.47% and 39.13% at D22 (Figure 2F).

Our results revealed that FAP^+^ cell ablation disturbed haematopoiesis in BM^5^ with a 70% reduction of Lin^−^Sca-1^+^ progenitors at D3, which recovered post-ablation at D22 (Figure 3A). Histological analysis revealed damaged marrow cavities at D3, whereas extracellular matrix deposits started filling the empty cavity with collagen and mucopolysaccharides at D7 (Figure 3B). The cell loss was also confirmed in thymus and spleen, 66% and 33% respectively (Figure 3C). In BM, the hematopoietic progenitor subsets, short-term (ST) HSCs were reduced by 68.9% whereas long-term (LT) did not (Figure 3D and supplemental figure 5), the oligopotent subsets (CLP, CMP, GMP) were reduced by 20-50% whilst MPP2 – whose genetic pathway is involved in homeostasis ^19, 20,21^ – increased by 50% but decreased 50% at D22 (Figures 3E and 3F). Moreover, MPP3 – a potent source of myeloid amplification in regenerative conditions - increased by 75% at D22 (Figure 3F) whilst the rest of the oligopotent subsets recovered, except CLP that maintained a similar percentage over the time. Results from extra medullar haematopoiesis activity in spleen showed reduced multipotent progenitors at D22 after DTX ablation whilst GMP, the oligopotent subset, increased by 150% (Figure 3G).

**Figure 3.**
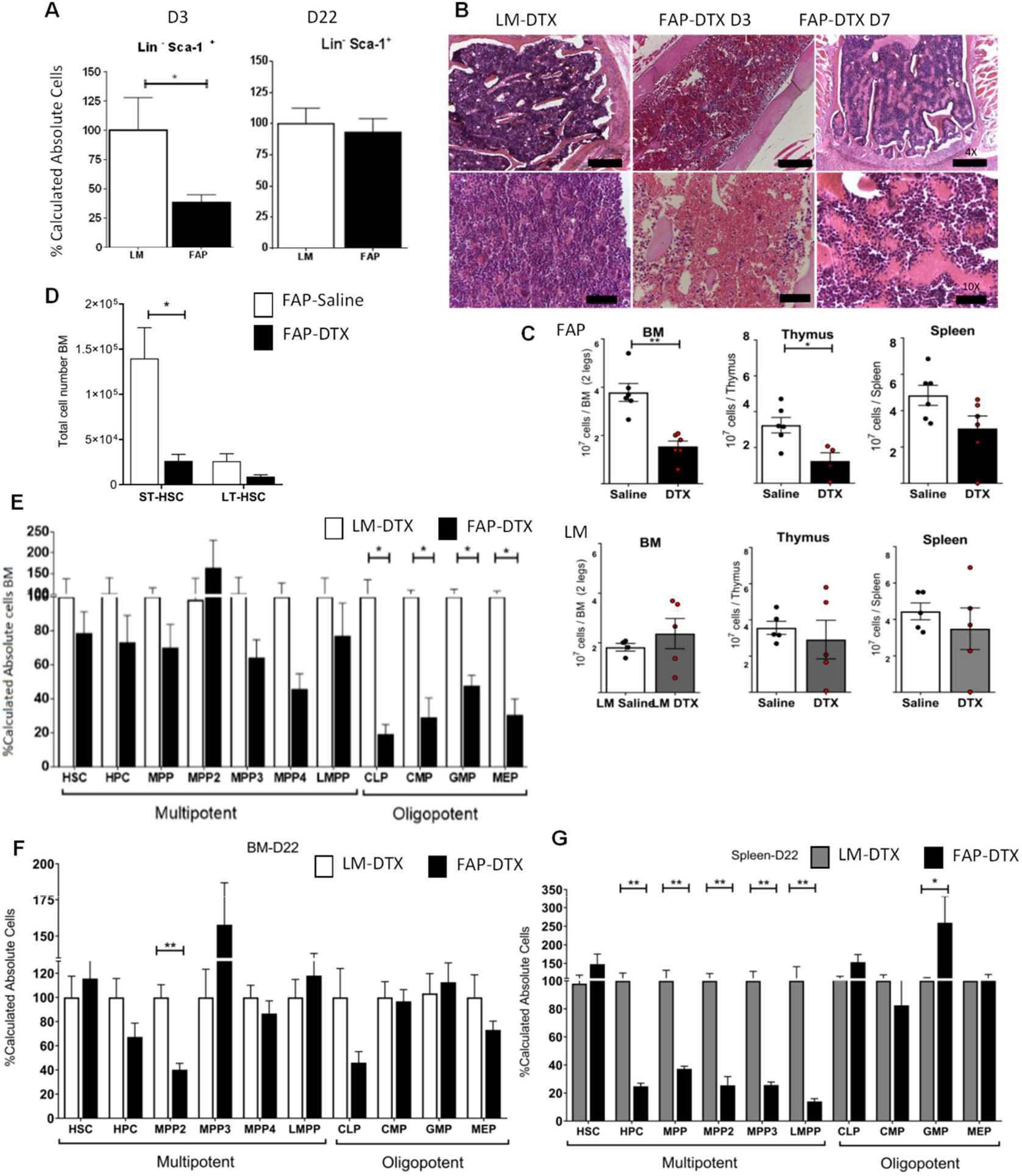
DTX-mediated FAP-cell ablation led to abnormal haematopoiesis in bone marrow and spleen. **(A)** Frequency of cell progenitor subset (Lin^−^Sca-1^+^) after cell ablation with DTX. n=6 per group, Mean ± SEM; α=0.05*,P<0.05. **(B)** Representative histological sections of the bone marrow cavity (Haematoxylin and eosin staining) after DTX administration. Scale bar represents 200μm. **(C)** Absolute cell count of nucleated cells from bone marrow, thymus and spleen at D3 after DTX administration. **(D)** Absolute cell count of short and long-term HSC (ST-HSC &LT-HSC) in bone marrow at D3 after DTX administration. **(E&F)** Frequency of the hematopoietic progenitors in bone marrow at **(E)** D3 and **(F)** D22 after DTX administration. **(G)** Frequency of the hematopoietic progenitors in the spleen at D22 after DTX administration. FAP saline and DTX (n=6 per group), LM saline and DTX (n=5 per group). Mean ± SEM, α=0.05 *,P<0.05; **,P<0.01.

### Syngeneic marrow stromal cells transplant ameliorates anaemia and rescues marrow progenitors

Immune competent mouse UBC-GFP was adopted as a donor of mesenchymal stromal cells (MSC) (Figure 4A) administering a single dose of 0.25×10^6^ (Low) and 0.5×10^6^ (High) 24 hours after the DTX regime (Figure 4B). BM nucleated cells were harvested and selected by adherence to plastic and expanded over four passages (Supplemental figures 9 and 10). The GFP expression (Supplemental figures 9 and 11) and cell surface markers, including FAP and Sca-1^+^ (Supplemental figure 10) confirmed progenitor stromal cell identity.

**Figure 4.**
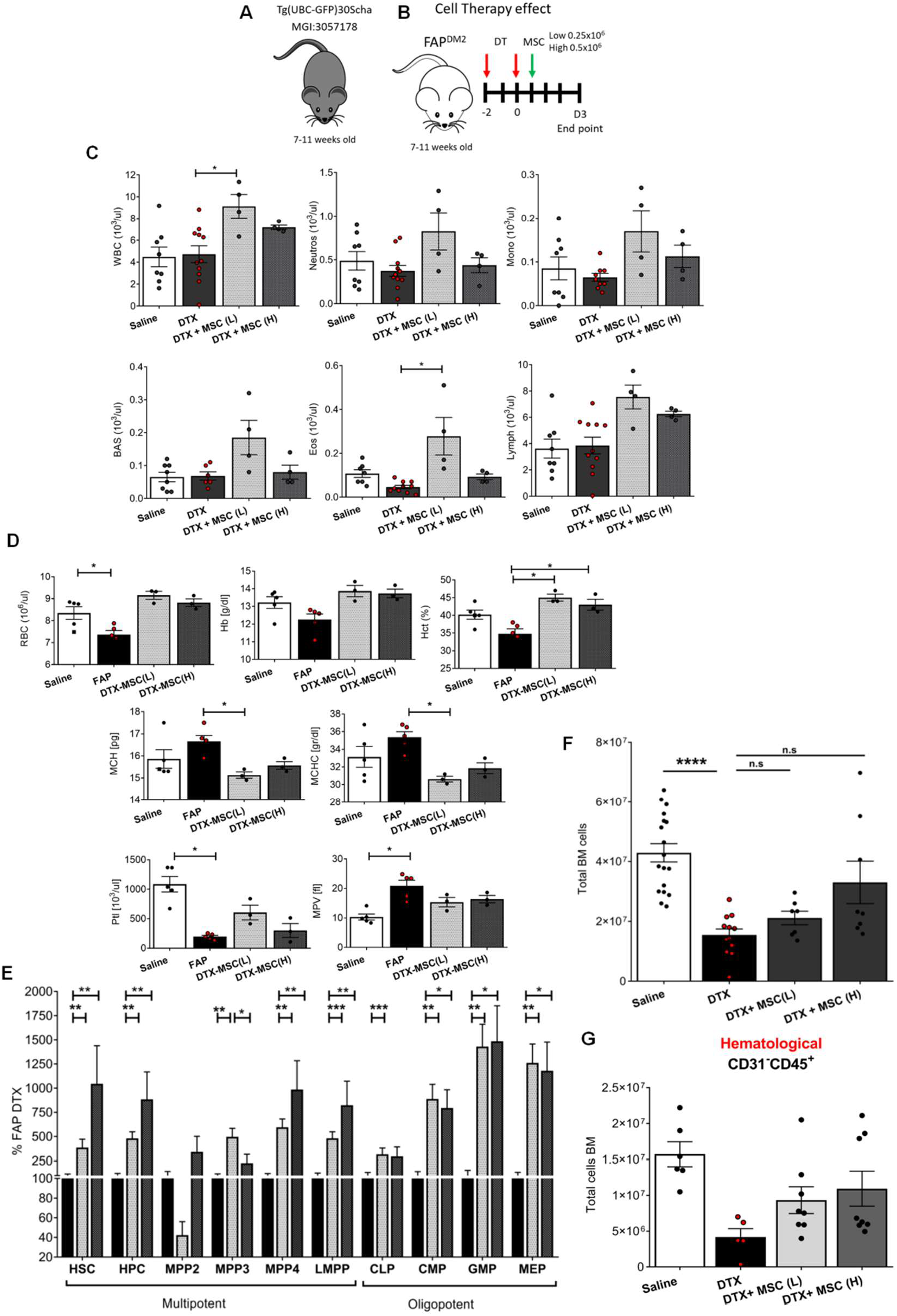
A single dose of MSCs restored anaemia and rescued progenitors in the marrow. The effect of the cell transplant was evaluated at D3 endpoint. **(A&B)** Schematic workflow of the adoptive transference of MSC from UBC-GFP mouse model in FAP^DM2^ mice after DTR ablation. **(C)** White blood parameters from peripheral blood. FAP-Saline (n=8), FAP-DTX (n=11), DTX+MSC (L and H) (n=4 per group). Mean ± SEM; α=0.05,*P<0.05; ***P<0.001. **(D)** Red blood parameters from peripheral blood. FAP-Saline and DTX (n=5 per group), DTX+MSC (L and H) (n=4 per group) Mean ± SEM; α=0.05*,P<0.05; ***P<0.001. **(E)** Frequency of the haematopoietic progenitor subsets in bone marrow. FAP-DTX (n=6), DTX+MSC (L and H) (n=8 per group). Mean±SEM, α=0.05, ****P<0.0001. **(F)** Absolute cell counts of nucleated bone marrow cells from processed femurs. FAP-Saline (n=22) and DTX (n=13), FAP-DTX+MSC (L+H) (n=7-8) Mean ± SEM, α=0.05, *P<0.05. **(G)** Absolute cell counts of haematological subset (CD45^+^ CD31^−^). FAP-Saline (n=4) and DTX (n=5), FAP-DTX+MSC (L+H) (n=7-8) Mean ± SEM, α=0.05, *P<0.05.

PB counts analysis at D3 after stromal cells transplant showed a 125% increase of WBC and in granulocytes subsets at a low dose, whilst the lymphocytes maintained similar levels when compared to FAP-DTX (Figure 4C). The total volume of blood (%Htc) increased by 30% at D3 post-transplant when compared to the FAP-DTX. The RBCs contained a similar amount of Hb (MCH, MCHC) whereas platelets quantity and their size reached the control (Figure 4D). Total marrow cellularity increased by 17% with a low dose of stromal cells, whereas it increased by 76% with the high dose (Figure 4F). Specifically, we identified that the haematological cluster (CD45^+^ CD31^−^) increased to 200% when compared to FAP-DTX ablation (Figure 4G). MSC transplant resulted in a significant impact on haematopoiesis, increasing the oligopotent GMP and MEP subsets fifteen fold, whilst CMP increased more than sevenfold.

Regarding the multipotent subsets, lymphoid-primed MPP4 increased tenfold versus fivefold with a high and low dose, respectively. Interestingly, MPP2 decreased after the low dose of the stromal cell was administered. (Figure 4E). Overall, MSC therapy boosted the cellular self-renewal in the marrow with a low dose of stromal cells restoring blood homeostasis and BM progenitor production.

## Discussion

This study demonstrated that (1) two doses of DTX ablated 50% of the BM (20×10^7^) in 72 hours with long-term effect in the organism homeostasis and, (2) that a single dose of ex vivo MSC (0.25-0.5 × 10^6^) administered i.v 24h after the DTX regime had a significant effect in ameliorating the damage produced by cells loss. FAP^+^ cell ablation led to a pathological phenotype affecting the PB counts (erythropenia, thrombocytopenia, and neutropenia), delayed lymphopenia and altered haematopoiesis with a significant reduction of progenitors in the marrow. Moreover, the liver enzymes (ALT and AST) responsible for cell removal within tissue damaged increased at D22. In such an emergency state of the BM, the extramedullary haematopoiesis in the spleen is activated whilst the thymus cellularity decreased as the lymphoid progenitor maturation is not required.

We show that FAP^+^ is not restricted to fibroblasts and overlaps with other stromal cell markers in osteolineage-chondrogenic spectrum confirming its role in bone homeostasis which is critical for HSC homing and BM transplantation involving a maturation-continuum from MSC transition^1^. Osteoblasts are involved in severe immune-mediated inflammatory diseases^18^, and recently confirmed the presence of FAP in osteolineage cells. Therefore, the physiological role of FAP^+^ stromal cells might be related to its contribution to bone homeostasis and its elimination set of all the alarms in 72 hours. In response to the damage, the MPP2 subset (lineage biased to erythrocytes and megakaryocytes) increased temporarily to maintain the homeostasis and ensure the oxygen to the tissues and the blood volume.

We suggest that transference of FAP^+^-MSC from the osteolineage and chondrogenic spectrum could contribute to alleviating the phenotype associated with immune-mediated inflammatory diseases, given its role in homeostasis and haematopoiesis within the BM. In this regards, FAP^DM2^ model presents a model first to eliminate FAP^+^ cells and then to explore the effect of specifically isolated stromal cells transference for severe immune-mediated inflammatory diseases.

## Conclusion

FAP^+^ cells represent a continuum of osteo-chondrocytes, and their depletion causes collateral damage to overlapping non-FAP^+^ stromal cells, endothelium and haematopoietic cells in the marrow and periphery. For the first time, we demonstrate that an intravenous infusion of a small dose of syngeneic marrow stromal cells can ameliorate the damage caused by FAP cell ablation. We demonstrate that stromal cell ablation models - such as FAP^DM2^ - can be used to deplete endogenous stromal cells and be used to explore the engraftment efficiency, function and therapeutic capacity of novel stromal cell subpopulations *in vivo.*

## Acknowledgements

The authors want to thank Ms Hemma Horan and Mr Patrick Costello for their technical assistance within the cell culture unit. The authors acknowledge the Veterinary hospital located in University College Dublin (UCD) for processing the blood. The authors wish to thank the Bio-Resources Unit technical, veterinary and administrative staff in NUI Galway for facilitating in-vivo studies and for their ongoing assistance, advice and support in animal husbandry, care and welfare. The authors acknowledge the facilities and technical assistance of the Flow Cytometry Core Facility at NUI Galway, which is funded, by NUI Galway, Science Foundation Ireland, and the Irish Government’s Programme for Research in Third Level Institutions, Cycle 5 and the European Regional Development Fund. The authors acknowledge the facilities and scientific and technical assistance of the Centre for Microscopy & Imaging at the National University of Ireland Galway (*www.imaging.nuigalway.ie).* The studies presented were supported by a European Union’s Seventh Framework Programme for research; technological development and demonstration under grant agreement n^o^ 315902. Marie Curie Initial Training Network DeCIDE (Decision-making within cells and differentiation entity therapies).

## Author contribution

SJE conceived and planned the study as well as revised the manuscript. ECR designed and performed the studies, found and implemented the technology, analysed and interpreted the data, prepared figures and wrote the manuscript. ECR with LW were responsible for the animal breeding, genotyping and, toxin/cells administration. LW, PL, SA helped in the tissue harvesting and sample processing. YD, LMD and LO contributed in the sample processing, analytical test and data interpretation. LRB contributed to the final version of the manuscript.

## References

1. Baryawno N, Przybylski D, Kowalczyk MS, Kfoury Y, Severe N, Gustafsson K et al. A Cellular Taxonomy of the Bone Marrow Stroma in Homeostasis and Leukemia. Cell 2019; 177(7):1915–1932.e1916. doi: 10.1016/j.cell.2019.04.040

2. Garin-Chesa P, Old LJ, Rettig WJ. Cell surface glycoprotein of reactive stromal fibroblasts as a potential antibody target in human epithelial cancers. Proceedings of the National Academy of Sciences 1990; 87(18):7235–7239. doi: 10.1073/pnas.87.18.7235

3. Bauer S, Jendro MC, Wadle A, Kleber S, Stenner F, Dinser R et al. Fibroblast activation protein is expressed by rheumatoid myofibroblast-like synoviocytes. Arthritis research & therapy 2006; 8(6):R171–R171. doi: 10.1186/ar2080

4. Kraman M, Bambrough PJ, Arnold JN, Roberts EW, Magiera L, Jones JO et al. Suppression of antitumor immunity by stromal cells expressing fibroblast activation protein-alpha. Science 2010; 330(6005):827–830. e-pub ahead of print 2010/11/06; doi: 10.1126/science.1195300

5. Roberts EW, Deonarine A, Jones JO, Denton AE, Feig C, Lyons SK et al. Depletion of stromal cells expressing fibroblast activation protein-alpha from skeletal muscle and bone marrow results in cachexia and anemia. J Exp Med 2013; 210(6):1137–1151. e-pub ahead of print 2013/05/29; doi: 10.1084/jem.20122344

6. Wosczyna MN, Konishi CT, Perez Carbajal EE, Wang TT, Walsh RA, Gan Q et al. Mesenchymal Stromal Cells Are Required for Regeneration and Homeostatic Maintenance of Skeletal Muscle. Cell Reports 2019; 27(7):2029–2035.e2025. doi: 10.1016/j.celrep.2019.04.074

7. Bae S, Park CW, Son HK, Ju HK, Paik D, Jeon C-J et al. Fibroblast activation protein α identifies mesenchymal stromal cells from human bone marrow. British Journal of Haematology 2008; 142(5):827–830. doi: 10.1111/j.1365-2141.2008.07241.x

8. Tran E, Chinnasamy D, Yu Z, Morgan RA, Lee CC, Restifo NP et al. Immune targeting of fibroblast activation protein triggers recognition of multipotent bone marrow stromal cells and cachexia. J Exp Med 2013; 210(6):1125–1135. e-pub ahead of print 2013/05/29; doi: 10.1084/jem.20130110

9. Decker M, Martinez-Morentin L, Wang G, Lee Y, Liu Q, Leslie J et al. Leptin-receptor-expressing bone marrow stromal cells are myofibroblasts in primary myelofibrosis. Nat Cell Biol 2017; 19(6):677–688. e-pub ahead of print 2017/05/10; doi: 10.1038/ncb3530

10. Zhou BO, Yue R, Murphy MM, Peyer JG, Morrison SJ. Leptin-receptor-expressing mesenchymal stromal cells represent the main source of bone formed by adult bone marrow. Cell Stem Cell 2014; 15(2):154–168. e-pub ahead of print 2014/06/24; doi: 10.1016/j.stem.2014.06.008

11. Wehmeyer C, Pap T, Buckley CD, Naylor AJ. The role of stromal cells in inflammatory bone loss. Clin Exp Immunol 2017; 189(1):1–11. e-pub ahead of print 2017/04/19; doi: 10.1111/cei.12979

12. Chung KM, Hsu SC, Chu YR, Lin MY, Jiaang WT, Chen RH et al. Fibroblast activation protein (FAP) is essential for the migration of bone marrow mesenchymal stem cells through RhoA activation. PLoS One 2014; 9(2):e88772. e-pub ahead of print 2014/02/20; doi: 10.1371/journal.pone.0088772

13. Santos AM, Jung J, Aziz N, Kissil JL, Pure E. Targeting fibroblast activation protein inhibits tumor stromagenesis and growth in mice. J Clin Invest 2009; 119(12):3613–3625. e-pub ahead of print 2009/11/19; doi: 10.1172/JCI38988

14. Zi F, He, J., He, D., Li, Y., Yang, L., & Cai, Z.. Fibroblast activation protein α in tumor microenvironment: Recent progression and implications. Molecular Medicine Reports 2015; 11:3203–3211. doi: https://doi.org/10.3892/mmr.2015.3197

15. Birbrair A, Frenette PS. Niche heterogeneity in the bone marrow. Ann N Y Acad Sci 2016; 1370(1):82–96. e-pub ahead of print 2016/03/26; doi: 10.1111/nyas.13016

16. Croft AP, Campos J, Jansen K, Turner JD, Marshall J, Attar M et al. Distinct fibroblast subsets drive inflammation and damage in arthritis. Nature 2019; 570(7760):246–251. doi: 10.1038/s41586-019-1263-7

17. Denton AE, Carr EJ, Magiera LP, Watts AJB, Fearon DT. Embryonic FAP^+^ lymphoid tissue organizer cells generate the reticular network of adult lymph nodes. The Journal of Experimental Medicine 2019: jem.20181705. doi: 10.1084/jem.20181705

18. Terashima A, Okamoto K, Nakashima T, Akira S, Ikuta K, Takayanagi H. Sepsis-Induced Osteoblast Ablation Causes Immunodeficiency. Immunity 2016; 44(6):1434–1443. e-pub ahead of print 2016/06/19; doi: 10.1016/j.immuni.2016.05.012

19. Pietras Eric M, Reynaud D, Kang Y-A, Carlin D, Calero-Nieto Fernando J, Leavitt Andrew D et al. Functionally Distinct Subsets of Lineage-Biased Multipotent Progenitors Control Blood Production in Normal and Regenerative Conditions. Cell stem cell 2015; 17(1):35–46. doi: http://dx.doi.org/10.1016/j.stem.2015.05.003

20. Pietras EM, Reynaud D, Kang YA, Carlin D, Calero-Nieto FJ, Leavitt AD et al. Functionally Distinct Subsets of Lineage-Biased Multipotent Progenitors Control Blood Production in Normal and Regenerative Conditions. Cell Stem Cell 2015; 17(1):35–46. e-pub ahead of print 2015/06/23; doi: 10.1016/j.stem.2015.05.003

21. Ceredig R. When one cell is enough. Stem cell research & therapy 2012; 3(1):1–1. doi: 10.1186/scrt92

