## Supplemental Information for "Bone marrow stromal cell transplantation ameliorates cytopenia caused by depletion FAP-α expressing cells"

**Supplemental Table 1. Transference of the DM2 transgene in both genders during the breeding process.**
**Supplemental Table 2. Hematopoietic stem cells subsets.**
**Supplemental Table 3. Cellular compartment phenotype in FAP-DTX ablation.**
**Supplemental Table 4. Cellular subsets phenotype in FAP-DTX ablation.**
**Supplemental Table 5. List of Antibodies**

**Supplemental Figure 1. Representative fap genotype by digesting ear biopsies.**
**Supplemental Figure 2. Cellular population complexity of BM-FAP$^{DM2}$.**
**Supplemental Figure 3. Molecular expression of *fap* in mouse bone marrow stroma.**
**Supplemental Figure 4. Molecular expression of stromal cell markers and factors in mouse bone marrow stroma.**
**Supplemental Figure 5. Flow cytometry gate strategy to identify the haematopoietic cell populations.**
**Supplemental Figure 6. Cell surface single proteins expression in BM after FAP-DTX ablation.**
**Supplemental Figure 7. Cellular compartments frequency in BM after FAP-DTX ablation.**
**Supplemental Figure 8. Comprehensive census of the cell subpopulations in BM after FAP-DTX ablation.**
**Supplemental Figure 9. Cell morphology of the expanded BM-UBC-GFP-MSCs**
**Supplemental Figure 10. Cell growth curve of the BM-UBC-GFP-MSCs**
**Supplemental Figure 11. Flow cytometry characterisation of the BM-UBC-GFP-MSCs.**

### Supplemental experiments procedures

**Mice:** $FAP^{DM2}$ mice provided to the National University of Ireland, Galway from Cambridge University whereas C57/Bl6-Tg (UBC-GFP) 30Scha/J (MGI: 3057178) purchased from Charles River. Mice breed in standard laboratory conditions of temperature (20 ± 2 °C), humidity (40–60%) and lighting (12:12 h light/dark cycle, lights on at 08:00h). FAP mice administered with saline or DTX *i.p* at 25ng/gr BW. Two injections performed at one day apart: day minus 2 and day zero for all experiments. A single dose of MSC ($0.250X10^6$ or $0.5X10^6$) in 100µl of saline administered *i.v* 24h after the second boost of DTX. All mice observed within the following 8h after any treatment given (saline, DTX either DTX+MSCs) following the animal care guidelines. Body weights recorded daily until the endpoint D3 or D22. The experimental procedures approved by the Animal Care and Research Ethics Committee of the National University of Ireland, Galway (NUIG) and carried out under the license from the Health Products Regulatory Authority (HPRA).

**Cardiocentesis:** To collect peripheral blood, the cardiac puncture performed immediately after euthanising to fill a 1ml syringe coated in advance with 10% 0.5M sterile EDTA for preventing clot formation. Blood kept at $4^oC$ until analysis no longer than 24hours post-collection. Serum obtained from blood collection without anticoagulant and settle until total coagulation (1-2h) at room temperature for enzymatic analysis. 250µl of clear serum transferred into a clean-labelled Eppendorf after centrifugation at 2500g for 20 min and stored at $-80^oC$ until analysis in UCD Veterinary Hospital.

**Genotyping:** DNA extracted from digesting ear biopsies using 100µl of lysis buffer ($N^o$402-E#Viagen) with 2µl ProteinaseK per mouse (0.3mg/ml). Low attachment tubes containing tissues with the lysis buffer incubated at 55°C for 3h and 45min at 700rpm in the thermoshaker. Tissue mix containing proteinase K heated at 85°C for 30 min and spun for 10 sec at 2000g to precipitate the hairs. Polymerase chain reaction (PCR) performed using ewr29 (Forward primer: GCATCCATGGAGAATGCAA 427 nucleotides into DTXR sequence) and ewr43 (Reverse primer: AATGGGAAGTCACGAAGGTG 3' primer) to magnify the DNA and evidence the presence of *fap*. PCR product analysed by electrophoresis in 1% agarose gel to evidence a band of 882 base pair bands.

**Tissue digestion:** Hind limbs bones from FAP and UBC-GFP mice processed to obtain total cell counts for cell analysis. The remaining flesh and the bone ends removed, bones rinsed with PBS1X. Bones crushed gently in the mortar using a pestle and α-MEN allowing to release the marrow which disassociated by pipetting up and down and kept in α–DMEN +20%FBS in the fridge. Chip bones digested with a buffer containing (α-MEM, collagenase P (0.2 mg/ml)(11213857001#Sigma-Aldrich), DNase I (0.1mg/ml)(DN25: 9003-98-9#Sigma-Aldrich), Dispase II (0.8mg/ml)(04942078001#Roche) 20 minutes at $37^oC$ in water bath. The supernatant containing the released cells from bone transferred into the tube containing the cells released at the crushing step. Digestion step performed 2-3 times until bones looked dusty. Total cell suspension filtered through 100µm cell strainer, spun at 400g 20 minutes. Erythrocytes in the pellet eliminated with 2ml of ACK Lysing Buffer for 2-3 min at RT Cell suspension counted for total cell number.

**Primary cell culture:** MSCs obtained from processed bones. Isolated nucleated cells from BM of UBC-GFP mice expanded under hypoxic cell culture conditions until passage 4 (37$^{o}$C and 5% $CO_2$ and 5% $O_2$). Nucleated cells free of erythrocytes platted at high density (1x$10^6$ cells/cm$^2$ in a flask T25), gently rinsed to eliminate the non-adherent cells with 1X PBS at 72h. Complete media (α-MEN+1% Penicillin/Streptomycin and 5%FBS) added every second day until 80-90% of confluence within 10 days. Trypsinized cells distributed for characterisation and functional tests and expanded by seeding at 5000 cells/cm$^2$ until they reached passage 4. When spare cells, they are frozen at 1x$10^6$/vial in CryoStor (C2874#Sigma Aldrich) and placed at -80$^{o}$C for cell banking.

**Flow cytometry:** BM, spleen, thymus cells from FAP in vivo experiments as well as the expanded BM-MSC from UBC-GP syngeneic mice stained and acquired in the BD FACS Canto II Cytometer. Analysis performed in FlowJo software.

Hematopoietic and endogenous stromal niches characterised by flow cytometric analysis. Depending on the experiment and test performed 1x$10^4$-$10^6$ cells used for each marker and the matched isotype or FMO control. FACS buffer (PBS1X +2% FBS+ 0.5mM EDTA) used for diluting antibodies. The process performed at 4$^{o}$C until cell acquisition. Viability dye eflour 780 (65-0865-18#eBioscience) used at the start of the staining or Sytox blue (S34857#ThermoFisher) previous acquisition at the cytometer.

<u>HSC subsets</u> (Supplemental Table 2) stained using a master mix tube contained Lineage (22-7770-72#BD), CD135 (FLT3) (12-1351#eBioscience), c-Kit (562417#BD), CD48 (560731#BD)/CD41 (133915#Medical supply), Sca-1(17-5981-81#eBioscience), and CD150 (562811#BD). CLP master mix tube contained exactly the same mixture as HSC tube but CD127 (IL-7R) (562959#BD) instead of CD150. MyP master mix tube contained Lineage, Sca-1(557405#BD), CD16/32 (553145#BD), CD34 (128607#Medical Supply), c-Kit (553356#BD) and CD127. FMOs tubes and other controls prepared.

Gating strategy for the HSC subsets: Morphological gate (FSC-A versus SSC-A) gated to eliminate the doublets (morphological- FSC-A versus FSC-H and complexity- SSC-A versus SSC-H eventually performed for high content acquisition. Viable cells e780-APC-Cy7 negative expression (633/780/60nm) versus FSC-A gated. Stem cell gate selected by Lineage-FITC$^-$(488/530/30nm) and Sca-1 APC (633/6620/30nm). Lin$^-$Sca-1$^+$ gated in a plot for c-kit- PE (488/585/42nm) versus Sca-1- APC (633/6620/30nm) for LS$^-$K (Lineage$^-$, Sca-1$^-$ and c-kit$^-$), LSK (Lineage$^-$ Sca-1$^+$ and c-kit$^+$) and LS$^{int}$K$^{int}$ (Lineage$^-$ Sca-1$^{int}$ and c-kit$^{int}$). LSK was gated for CD48/CD41 PE-Cy7 (488/780/60nm) and CD150-VB421 (405/450/50nm) to examine HPC (CD48$^+$CD150$^-$), MPP (CD48$^-$CD150$^-$) and HSC (CD48$^-$ CD150$^+$) populations. HPC population gated to examine under CD135-PE (488/585/42) and CD150-VB421 (405/450/50 nm), LMPP (CD135$^+$CD150$^{-/+}$). CLP population gated from LS$^{int}$K$^{int}$ (Lineage$^-$ Sca-1$^{int}$ and c-kit$^{int}$) population gated for positive expression of IL-7R-VB421 (405/450/50nm). Oligopotent populations (GMP, CMP and MEP) gated in a plot of Lineage$^-$, Sca-1$^-$ FITC (488/530/30nm) versus FSC-A. Negative population gated for IL-7R-VB421 (405/450/50nm) versus FSC-A to select the negative fraction and plot for c-kit-APC (633/6620/30nm). The positive fraction gated in a plot for CD13/32-PE (488/585/42nm) versus CD34 PerCP-Cy5.5 (488/670LPnm) to define MEP (CD16/32$^{-}$ CD34$^{-}$), CMP (CD16/32$^{int}$CD34$^{int}$) and GMP (CD16/32$^{+}$CD34$^{+}$). MPP2 (FLT3$^{-}$ CD150$^{+}$CD48$^{+}$), MPP3 (FLT3$^{-}$, CD150$^{-}$CD48$^{+}$), HSC-LT (FLT3$^{-}$CD150$^{+}$CD48$^{-}$) and HSC-ST (FLT3$^{-}$CD150$^{-}$CD48$^{-}$) gated from LSK pool created using Lineage-FITC (488/530/30 nm) versus FSC-A, Lin- plot for c-kit-APC and Sca-1(633/6620/30nm) versus Lineage-FITC (488/530/30nm). The double-positive population LSK gated for FLT3$^{-}$CD135-PE (488/585/42). The two gates based on the positive and negative expression of it plotted for CD150-VB421 (405/450/50 nm) versus CD48-PE-Cy7 (488/780/60). MPP4 (FLT3$^{+}$CD150$^{-}$CD48$^{+}$) gated from FLT3$^{+}$ fraction.

For endogenous stromal cells analysis, Ter119 (557915#BD), CD31 (102508#eBioscience) and CD45 (553080#BD) used to evaluate FAP expression (AF3715#R&D), Syndecan 2 (CD362) (FAB2965A#R&D), Platelet-derived growth factor receptor alpha (PDGFR-$\alpha$) (135910#BD, 135910#Medical supply), Leptin-R (BAF497#R&D) and gp38 (Podoplanin) (12-5381-82#eBioscience). Cell receptors blocked CD16/32 (553142#BD) 15min, and without washing, FAP ab added for 15 min. In a second staining step, cell receptors blocked again, and the biotinylated secondary Ab (B3148#Sigma) added. Streptavidin used upon rinsed and incubated for 30 min. Expanded plastic adherent from UBC-GFP mice stained with FAP, SDC2, (PDGFR–$\alpha$), gp38, Sca-1, CD31(102417#MedicalSupply), CD73 (550741#BD), CD54 (12054181#Medical Supply) and MHC-II (553570#BD), CD11b(557657#BD), F4/80 (123116#Medical supply) and CD86(560582#BD) for cell characterisation. CaliBRITE unlabelled beads (01-2222-42#BD) used to calculate the total cell number of cells within each cell population in tissue/mouse. Bead solution prepared by adding 40 drops to 10ml of FACS buffer. Beads count in the haemocytometer to calculate the total number of cells within the cell subset of interest. A volume of 50µl of beads suspension added to the stained cell tube just before cytometer acquisition.

The compensation matrix created by adding 1 drop of UltraComp eBeads (01-2222-42#ThermoFisher) per fluorophore. Incubated 10minutes, washed with 2ml of FACS buffer, centrifuged 1500rpm 5 min and left in 200µl of FACS buffer. Brilliant buffer required to reduce the non-specific interactions when V421 and V510 fluorophores added in the same test. The resulting counts from samples referred to the average of the control group. The average of the control values was referred to 100%, and values from the study group referred to this percentage.

**Visualisation and clustering.** The dimensionality of the 1x10$^{6}$ cell dataset reduced to project 200,000 events by using t-Distributed stochastic neighbour embedding (t-SNE) with FlowJo software. Aligned canonical correlation analysis used as a basis for partitioning the dataset into clusters using a shared nearest neighbour modularity optimisation algorithm. Based on the expression levels of the four main clusters (CD45$^{+}$CD31$^{-}$, CD45$^{+}$CD3$^{+}$, CD45$^{-}$CD31$^{+}$ and CD45$^{-}$CD31$^{-}$), the four-cell compartments displayed distinct visual clusters. The same algorithm used to extend the analysis, including the stromal cell markers within each cluster.

**Histology**: Bone decalcification and paraffin-embedded processes performed before sectioning. Bones fixed in 10% formalin for 24h. When decalcification required, bones transferred into Kristensen's Solution (Sodium fumarate and Formic acid-F1506 and 399388#Sigma) rinsed with running tap water before preserved with 30% sucrose (S9378#Sigma) solution at 4$^{o}$C until the bones sank. When paraffin-embedded tissues, automatic tissue processor used upon 10% formalin (HT501128#Sigma) fixation. Tissue cassettes placed into 70% ethanol for at least 16h to start the dehydration process (increasing percentages of ethanol, followed by 3 cycles of xylene (534056#Sigma) and paraffin wax in the tissue processor (ASP300# Leica). A manual rotary microtome (RM2235#Leica) used for paraffin-embedding method whereas manual rotary cryostat (CM1850#Leica) used for cryopreserved tissues. All tissues sectioned <10μm thick. **H&E staining**: Sections stained with Mayer's Haematoxylin (51275#Sigma) (3 min) and Eosin dye (CI 45380, A0822,0100 #Lennox) (1.5 min). To dehydrate the tissues a series of increased graded ethanol solutions (50, 70, 95, and 2 changes of 100%) used (10 seconds for the 50, 75 and 90 and 2 min for each change at 100%). Samples cleared with xylene (2 changes of 10 min each). Sections covered with DPX mounting medium and a coverslip applied to preserve the staining. Slices kept in the fume hood overnight and transferred into the oven at 37$^{o}$C for at least 16h before examination in the bright field microscope (Leica DM2500).

**Statistical Analysis:** Statistics analysed using GraphPad Prism software and graphs generated using both GraphPad Prism and Microsoft Excel when required. Non-parametric analysis, two-tailed Mann Whitney and Kruskal-Wallis tests used for studies with two or more than two groups, respectively. One-way ANOVA by Tukey's post hoc test used. When two-ways ANOVA Sidak's multiple comparison test used. Statistical significance for $\alpha=0.05$ denoted as follows $*p\leq0.05$, $**p\leq0.01$, $***p\leq0.001$

### Supplemental Tables

| | | Genotype | | | | | | | fap+ | |
|---|---|---|---|---|---|---|---|---|---|---|
| | | Male (♂) | | | Female (♀) | | | | | |
| Genotype number | number of mice | number of mice | *fap+* | LM | number of mice | *fap+* | LM | number of mice | **Number (♂+♀)** | **% (♂+♀)** |
| **1** | 14 | **9** | 3 | 6 | **5** | 1 | 4 | **14** | 4 | **28.57** |
| **2** | 21 | **13** | 5 | 8 | **8** | 3 | 5 | **21** | 8 | **38.10** |
| **3** | 17 | **10** | 4 | 6 | **7** | 4 | 3 | **17** | 8 | **47.06** |
| **4** | 28 | **14** | 7 | 7 | **14** | 7 | 7 | **28** | 14 | **50.00** |
| **5** | 31 | **14** | 9 | 5 | **17** | 11 | 6 | **31** | 20 | **64.52** |
| **6** | 34 | **19** | 8 | 11 | **13** | 10 | 3 | **34** | 18 | **52.94** |
| **7** | 24 | **14** | 10 | 4 | **10** | 6 | 4 | **24** | 16 | **66.67** |
| **8** | 13 | **13** | 2 | 1 | **10** | 6 | 4 | **13** | 8 | **61.54** |
| **9** | 47 | **20** | 5 | 15 | **23** | 12 | 11 | **47** | 17 | **36.17** |
| **10** | 25 | **12** | 8 | 4 | **13** | 6 | 7 | **25** | 14 | **56.00** |
| **11** | 112 | **54** | 33 | 21 | **58** | 33 | 25 | **112** | 66 | **58.93** |
| **12** | 17 | **10** | 5 | 5 | **6** | 2 | 4 | **17** | 7 | **41.18** |

| | **mice numbers** | | |
|---|---|---|---|
| | Total | ***fap+*** | ***LM*** |
| | 383 | 200 | 176 |
| ♂ | 202 | 99 | 93 |
| ♀ | 184 | 101 | 83 |

| | **Percentage (%)** | | |
|---|---|---|---|
| | Total | ***fap+*** | ***LM*** |
| ♂ | 52.7 | 49 | 46.03 |
| ♀ | 48 | 53.8 | 45.1 |

**Supplemental Table 1. Transference of the DM2 transgene in both genders during the breeding process.**
The table shows the number of genotyped mice together with the number and percentage of gene transference obtained from the twelve breedings. The small table summarises the results by gender and condition expressed as a percentage and total numbers.

| | Cell surface combination markers | Short name | Mame |
|---|---|---|---|
| Progenitors cells | Lin$^-$ Sca-1$^+$ | Lin$^-$Sca$^+$ | Progenitors cells |
| Progenitors Intermediares | Lin$^-$ Sca-1$^-$ c-kit$^+$ | LS$^-$K$^+$ | Intermediate population 1 |
| | Lin$^-$ Sca-1$^{int}$ c-kit$^{int}$ | LS$^{int}$ K$^{int}$ | Intermediate population 2 |
| | Lin$^-$ Sca-1$^+$ c-kit$^+$ | LSK | Intermediate population 3 |
| Multipotent | LSK (C48$^-$CD150$^+$) | HSC | Hematopoietic Stem Cell |
| | LSK (CD48$^+$ CD150$^-$) | HPC | Hematopoietic Progenitor Cell |
| | LSK (C48$^-$CD150$^-$) | MPP | Multipotent pogenitor |
| | LSK (Flt3$^-$ CD48$^+$ CD150$^-$) | MPP3 | Multipotent pogenitor 3 |
| | LSK (Flt3$^-$ CD48$^+$ **CD150$^+$**) | MPP2 | Multipotent pogenitor 2 |
| | LSK ( **Flt3$^+$** CD48$^+$ CD150$^-$) | MPP4 | Multipotent pogenitor 4 |
| | LSK (Flt3$^+$ CD48$^+$ **CD150$^+$**) | LMPP | Lymphoid multipotential progenitors |
| Oligopotent | LS$^-$K$^+$ (IL-7R$^-$ CD16/32$^{lo}$ CD34$^{hi}$) | CMP | Common myeloid progenitors |
| | LS$^{int}$ K$^{int}$ ( IL-7R$^+$) | CLP | Common lymphoid progenitors |
| | LS$^-$K$^+$ (IL-7R$^-$ CD16/32$^{hi}$ CD34$^{hi}$) | GMP | Granulocyte-macrophage progenitors |
| | LS$^-$K$^+$ (IL-7R$^-$ CD16/32$^{lo}$ CD34$^{lo}$) | MEP | Megakaryocyte–erythroid progenitor cell |

**Supplemental Table 2. Hematopoietic stem cells subsets.**
Description of the analysed subsets based on the cell surface markers.

| A | | Absolute number of cells in BM | | % cells in BM | | |
|---|---|---|---|---|---|---|
| | | Mean | SEM | Mean | SEM | % Ablation |
| CD45$^+$CD31$^-$ | LM Saline | 2.34E+07 | 2.05E+06 | 100.00 | 13.81 | 37.87 |
| | LM DTX | 2.34E+07 | 2.05E+06 | 62.13 | 5.46 | |
| CD45+CD31+ | LM | 2.59E+06 | 6.22E+04 | 100.00 | 2.42 | 13.04 |
| | LM DTX | 2.25E+06 | 2.70E+05 | 86.96 | 10.43 | |
| CD45-CD31+ | LM | 1.23E+05 | 3.50E+04 | 100.00 | 28.54 | 60.64 |
| | LM DTX | 4.83E+04 | 1.17E+04 | 39.37 | 9.56 | |
| CD45-CD31- | LM | 2.45E+06 | 2.85E+05 | 100.00 | 11.59 | 19.47 |
| | LM DTX | 1.97E+06 | 1.94E+05 | 80.53 | 7.94 | |

| B | | Absolute number of cells in BM | | % cells in BM | | |
|---|---|---|---|---|---|---|
| | | Mean | SEM | Mean | SEM | % Ablation |
| CD45$^+$CD31$^-$ | FAP Saline | 2.96E+07 | 4.47E+06 | 100.00 | 15.09 | 58.67 |
| | FAP DTX | 1.22E+07 | 1.34E+06 | 41.33 | 4.48 | |
| CD45+CD31+ | FAP Saline | 2.23E+06 | 2.16E+05 | 100.00 | 9.79 | 30.34 |
| | FAP DTX | 1.56E+06 | 1.38E+05 | 69.66 | 6.18 | |
| CD45-CD31+ | FAP Saline | 3.20E+05 | 1.66E+05 | 100.00 | 41.45 | 75.37 |
| | FAP DTX | 5.42E+04 | 1.61E+04 | 24.63 | 7.32 | |
| CD45-CD31- | FAP Saline | 2.17E+06 | 6.65E+05 | 100.00 | 32.67 | 82.55 |
| | FAP DTX | 4.36E+05 | 1.62E+05 | 17.45 | 6.47 | |

**Supplemental Table 3. Cellular compartment phenotype in FAP-DTX ablation.**
Data collection contains the absolute cell counts, the frequency and the percentage of ablation for each coloured-code cellular compartments in (A) LM and (B) FAP transgenic mice at D3 after DTX administration or saline as a control. n=4 Mean ± SEM.

| | | Absolute number of cells in BM | | | | |
|---|---|---|---|---|---|---|
| | | Mean | SEM | | Mean | SEM |
| CD45$^+$CD31$^-$ | LM Saline | 2.34E+07 | 2.05E+06 | FAP Saline | 2.96E+07 | 4.47E+06 |
| | LM DTX | 2.34E+07 | 2.05E+06 | FAP DTX | 1.22E+07 | 1.34E+06 |
| FAP-a$^+$ | LM Saline | 1.99E+06 | 4.57E+04 | FAP Saline | 2.63E+06 | 2.83E+05 |
| | LM DTX | 1.25E+06 | 2.70E+05 | FAP DTX | 1.12E+06 | 8.99E+04 |
| gp38$^+$ | LM Saline | 9.23E+03 | 8.53E+02 | FAP Saline | 9.17E+03 | 1.32E+03 |
| | LM DTX | 7.16E+03 | 1.09E+03 | FAP DTX | 5.75E+03 | 5.80E+02 |
| Leptin-R$^+$ | LM Saline | 8.68E+04 | 5.32E+03 | FAP Saline | 8.27E+04 | 1.15E+04 |
| | LM DTX | 7.08E+04 | 9.18E+03 | FAP DTX | 7.16E+04 | 1.03E+04 |
| PDFGR-a$^+$ | LM Saline | 4.24E+04 | 3.29E+03 | FAP Saline | 3.80E+04 | 5.55E+03 |
| | LM DTX | 2.82E+04 | 4.79E+03 | FAP DTX | 2.28E+04 | 2.45E+03 |
| SDC2$^+$ | LM Saline | 1.63E+05 | 4.52E+04 | FAP Saline | 1.19E+05 | 2.04E+04 |
| | LM DTX | 7.91E+04 | 8.86E+03 | FAP DTX | 4.68E+04 | 6.32E+03 |

| | | % cells in BM | | | | |
|---|---|---|---|---|---|---|
| | | Mean | SEM | | Mean | SEM |
| CD45$^+$CD31$^-$ | LM Saline | 100.00 | 13.81 | FAP Saline | 100.00 | 15.09 |
| | LM DTX | 62.13 | 5.46 | FAP DTX | 41.33 | 4.48 |
| FAP-a$^+$ | LM Saline | 5.29 | 0.12 | FAP Saline | 8.88 | 0.95 |
| | LM DTX | 3.33 | 0.72 | FAP DTX | 3.80 | 0.31 |
| gp38$^+$ | LM Saline | 0.03 | 0.00 | FAP Saline | 0.03 | 0.00 |
| | LM DTX | 0.02 | 0.00 | FAP DTX | 0.02 | 0.00 |
| Leptin-R$^+$ | LM Saline | 0.23 | 0.01 | FAP Saline | 0.28 | 0.04 |
| | LM DTX | 0.19 | 0.02 | FAP DTX | 0.24 | 0.04 |
| PDFGR-a$^+$ | LM Saline | 0.11 | 0.01 | FAP Saline | 0.13 | 0.02 |
| | LM DTX | 0.07 | 0.01 | FAP DTX | 0.08 | 0.01 |
| SDC2$^+$ | LM Saline | 0.44 | 0.12 | FAP Saline | 0.40 | 0.07 |
| | LM DTX | 0.21 | 0.02 | FAP DTX | 0.16 | 0.02 |

| % Ablation | LM | FAP |
|---|---|---|
| CD45$^+$CD31$^-$ | 37.87 | 58.67 |
| FAP-a$^+$ | 37.07 | 57.21 |
| gp38$^+$ | 30.00 | 33.33 |
| Leptin-R$^+$ | 19.35 | 13.39 |
| PDFGR-a$^+$ | 35.56 | 41.18 |
| SDC2$^+$ | 51.72 | 60.87 |

| | | Absolute number of cells in BM | | | | |
|---|---|---|---|---|---|---|
| | | Mean | SEM | | Mean | SEM |
| CD45$^+$CD31$^+$ | LM Saline | 2.59E+06 | 6.22E+04 | FAP Saline | 2.23E+06 | 2.16E+05 |
| | LM DTX | 2.25E+06 | 2.70E+05 | FAP DTX | 1.56E+06 | 1.38E+05 |
| FAP-a$^+$ | LM Saline | 5.23E+04 | 2.67E+03 | FAP Saline | 5.00E+04 | 6.88E+03 |
| | LM DTX | 4.09E+04 | 9.95E+03 | FAP DTX | 3.90E+04 | 3.84E+03 |
| gp38$^+$ | LM Saline | 3.43E+04 | 2.78E+03 | FAP Saline | 2.25E+04 | 4.10E+03 |
| | LM DTX | 1.46E+04 | 2.86E+03 | FAP DTX | 1.76E+04 | 2.10E+03 |
| Leptin-R$^+$ | LM Saline | 1.02E+04 | 5.98E+02 | FAP Saline | 8.87E+03 | 2.12E+03 |
| | LM DTX | 9.13E+03 | 1.53E+03 | FAP DTX | 1.42E+04 | 1.88E+03 |
| PDFGR-a$^+$ | LM Saline | 1.44E+04 | 8.91E+02 | FAP Saline | 9.55E+03 | 1.52E+03 |
| | LM DTX | 7.90E+03 | 1.40E+03 | FAP DTX | 1.05E+04 | 7.12E+02 |
| SDC2$^+$ | LM Saline | 1.45E+05 | 1.30E+04 | FAP Saline | 1.02E+05 | 9.62E+03 |
| | LM DTX | 8.97E+04 | 2.05E+04 | FAP DTX | 6.21E+04 | 4.72E+03 |

| | | % cells in BM | | | | |
|---|---|---|---|---|---|---|
| | | Mean | SEM | | Mean | SEM |
| CD45$^+$CD31$^+$ | LM Saline | 100.00 | 2.42 | FAP Saline | 100.00 | 9.79 |
| | LM DTX | 86.96 | 10.43 | FAP DTX | 69.66 | 6.18 |
| FAP-a$^+$ | LM Saline | 2.02 | 0.10 | FAP Saline | 2.24 | 0.31 |
| | LM DTX | 1.58 | 0.38 | FAP DTX | 1.75 | 0.17 |
| gp38$^+$ | LM Saline | 1.33 | 0.11 | FAP Saline | 1.01 | 0.18 |
| | LM DTX | 0.56 | 0.11 | FAP DTX | 0.79 | 0.09 |
| Leptin-R$^+$ | LM Saline | 0.40 | 0.02 | FAP Saline | 0.40 | 0.09 |
| | LM DTX | 0.35 | 0.06 | FAP DTX | 0.64 | 0.08 |
| PDFGR-a$^+$ | LM Saline | 0.55 | 0.03 | FAP Saline | 0.43 | 0.07 |
| | LM DTX | 0.31 | 0.06 | FAP DTX | 0.47 | 0.03 |
| SDC2$^+$ | LM Saline | 5.61 | 0.51 | FAP Saline | 4.56 | 0.43 |
| | LM DTX | 3.47 | 0.79 | FAP DTX | 2.78 | 0.21 |

| % Ablation | LM | FAP |
|---|---|---|
| CD45$^+$CD31$^+$ | 13.04 | 30.34 |
| FAP-a$^+$ | 21.56 | 22.19 |
| gp38$^+$ | 57.55 | 21.78 |
| Leptin-R$^+$ | 10.76 | 61.01 |
| PDFGR-a$^+$ | 44.80 | 9.94 |
| SDC2$^+$ | 38.18 | 39.07 |

| | | Absolute number of cells in BM | | | | |
|---|---|---|---|---|---|---|
| | | Mean | SEM | | Mean | SEM |
| CD45$^-$CD31$^-$ | LM Saline | 2.45E+06 | 2.85E+05 | FAP Saline | 2.17E+06 | 6.65E+05 |
| | LM DTX | 1.97E+06 | 1.94E+05 | FAP DTX | 4.36E+05 | 1.62E+05 |
| FAP-a$^+$ | LM Saline | 1.85E+04 | 3.27E+03 | FAP Saline | 1.65E+04 | 3.63E+03 |
| | LM DTX | 7.16E+03 | 1.86E+03 | FAP DTX | 2.73E+03 | 5.14E+02 |
| gp38$^+$ | LM Saline | 9.04E+03 | 8.36E+02 | FAP Saline | 9.96E+03 | 3.11E+03 |
| | LM DTX | 4.30E+03 | 4.15E+02 | FAP DTX | 1.50E+03 | 3.02E+02 |
| Leptin-R$^+$ | LM Saline | 1.38E+04 | 3.11E+03 | FAP Saline | 1.94E+04 | 7.45E+03 |
| | LM DTX | 5.89E+03 | 1.53E+03 | FAP DTX | 1.93E+03 | 3.80E+02 |
| PDFGR-a$^+$ | LM Saline | 3.70E+03 | 4.74E+02 | FAP Saline | 5.10E+03 | 2.11E+03 |
| | LM DTX | 2.23E+03 | 5.08E+02 | FAP DTX | 6.66E+02 | 1.43E+02 |
| SDC2$^+$ | LM Saline | 2.97E+04 | 5.66E+03 | FAP Saline | 3.03E+04 | 6.89E+03 |
| | LM DTX | 1.61E+04 | 1.16E+03 | FAP DTX | 3.73E+03 | 7.93E+02 |

| | | % cells in BM | | | | |
|---|---|---|---|---|---|---|
| | | Mean | SEM | | Mean | SEM |
| CD45$^-$CD31$^-$ | LM Saline | 100.00 | 11.59 | FAP Saline | 100.00 | 32.67 |
| | LM DTX | 80.53 | 7.94 | FAP DTX | 17.45 | 6.47 |
| FAP-a$^+$ | LM Saline | 0.76 | 0.13 | FAP Saline | 0.66 | 0.15 |
| | LM DTX | 0.29 | 0.08 | FAP DTX | 0.11 | 0.02 |
| gp38$^+$ | LM Saline | 0.37 | 0.03 | FAP Saline | 0.40 | 0.12 |
| | LM DTX | 0.18 | 0.02 | FAP DTX | 0.06 | 0.01 |
| Leptin-R$^+$ | LM Saline | 0.57 | 0.13 | FAP Saline | 0.78 | 0.30 |
| | LM DTX | 0.24 | 0.06 | FAP DTX | 0.08 | 0.02 |
| PDFGR-a$^+$ | LM Saline | 0.15 | 0.02 | FAP Saline | 0.21 | 0.08 |
| | LM DTX | 0.09 | 0.02 | FAP DTX | 0.03 | 0.01 |
| SDC2$^+$ | LM Saline | 1.22 | 0.23 | FAP Saline | 1.22 | 0.28 |
| | LM DTX | 0.66 | 0.05 | FAP DTX | 0.15 | 0.03 |

| % Ablation | LM | FAP |
|---|---|---|
| CD45$^-$CD31$^-$ | 19.47 | 82.55 |
| FAP-a$^+$ | 61.72 | 83.33 |
| gp38$^+$ | 52.70 | 85.00 |
| Leptin-R$^+$ | 57.71 | 90.03 |
| PDFGR-a$^+$ | 39.34 | 87.80 |
| SDC2$^+$ | 45.68 | 87.65 |

| | | Absolute number of cells in BM | | | | |
|---|---|---|---|---|---|---|
| | | Mean | SEM | | Mean | SEM |
| CD45$^-$CD31$^+$ | LM Saline | 1.23E+05 | 3.50E+04 | FAP Saline | 3.20E+05 | 1.66E+05 |
| | LM DTX | 4.83E+04 | 1.17E+04 | FAP DTX | 5.42E+04 | 1.61E+04 |
| FAP-a$^+$ | LM Saline | 5.50E+03 | 4.01E+02 | FAP Saline | 1.95E+04 | 9.48E+03 |
| | LM DTX | 3.60E+03 | 1.70E+02 | FAP DTX | 2.11E+03 | 6.27E+02 |
| gp38$^+$ | LM Saline | 2.10E+03 | 2.03E+02 | FAP Saline | 1.25E+03 | 2.04E+02 |
| | LM DTX | 1.32E+03 | 3.79E+02 | FAP DTX | 2.73E+02 | 5.26E+01 |
| Leptin-R$^+$ | LM Saline | 1.74E+02 | 4.11E+01 | FAP Saline | 2.80E+02 | 5.61E+01 |
| | LM DTX | 1.99E+02 | 5.40E+01 | FAP DTX | 7.10E+01 | 2.84E+01 |
| PDFGR-a$^+$ | LM Saline | 2.71E+02 | 6.86E+01 | FAP Saline | 2.71E+02 | 3.10E+01 |
| | LM DTX | 8.88E+01 | 1.44E+01 | FAP DTX | 7.29E+01 | 1.27E+01 |
| SDC2$^+$ | LM Saline | 7.77E+03 | 5.78E+02 | FAP Saline | 7.85E+03 | 1.10E+03 |
| | LM DTX | 4.83E+03 | 9.69E+02 | FAP DTX | 1.92E+03 | 1.94E+02 |

| | | % cells in BM | | | | |
|---|---|---|---|---|---|---|
| | | Mean | SEM | | Mean | SEM |
| CD45$^-$CD31$^+$ | LM Saline | 100.00 | 28.54 | FAP Saline | 100.00 | 41.45 |
| | LM DTX | 39.37 | 9.56 | FAP DTX | 24.63 | 7.32 |
| FAP-a$^+$ | LM Saline | 4.49 | 0.33 | FAP Saline | 3.17 | 0.75 |
| | LM DTX | 2.93 | 0.14 | FAP DTX | 0.96 | 0.29 |
| gp38$^+$ | LM Saline | 1.71 | 0.17 | FAP Saline | 0.57 | 0.09 |
| | LM DTX | 1.08 | 0.31 | FAP DTX | 0.12 | 0.02 |
| Leptin-R$^+$ | LM Saline | 0.14 | 0.03 | FAP Saline | 0.13 | 0.03 |
| | LM DTX | 0.16 | 0.04 | FAP DTX | 0.05 | 0.01 |
| PDFGR-a$^+$ | LM Saline | 0.22 | 0.06 | FAP Saline | 0.12 | 0.01 |
| | LM DTX | 0.07 | 0.01 | FAP DTX | 0.03 | 0.00 |
| SDC2$^+$ | LM Saline | 6.34 | 0.47 | FAP Saline | 3.56 | 0.50 |
| | LM DTX | 3.94 | 0.79 | FAP DTX | 0.87 | 0.09 |

| % Ablation | LM | FAP |
|---|---|---|
| CD45$^-$CD31$^+$ | 60.64 | 75.37 |
| FAP-a$^+$ | 34.69 | 69.67 |
| gp38$^+$ | 36.84 | 78.41 |
| Leptin-R$^+$ | 16.07 | 62.00 |
| PDFGR-a$^+$ | 68.18 | 73.47 |
| SDC2$^+$ | 37.81 | 75.47 |

**Supplemental Table 4. Cellular subsets phenotype in FAP-DTX ablation.**
Data collection of the cellular subsets expressing FAP, gp38, LeptinR, PDFGR$\alpha$ and SDC2 within each coloured-code cellular compartment in BM. Each section contains a table with the absolute cell counts, another table with the calculated frequency and the last table contains the calculated percentage of ablation for each cell sub-population analysed in both, LM and FAP at D3 after DTX administration or saline as a control. n=4 mean ± SEM.

| Ab | Clone | Isotype | Label | Dilution | Supplier code | Manufacturer supplier |
|---|---|---|---|---|---|---|
| Flow Cytometry | | | | | | |
| FAP-α | | Sheep IgG | Purified | 1/50 | AF3715 | Bio-teche(R&D systems),UK |
| CD16/32 | | Rat IgG2b$_{,k}$ | Purified | 1/100 | 553142 | BD, UK |
| α-Sheep | GT-34 | IgG1 | Biotin | 1/1000 | B3148 | Sigma, UK |
| Leptin-R | | Goat IgG1 | Biotin | 1/200 | BAF 497 | Bio-teche(R&D systems),UK |
| PDFGR-α | APA5 | Rat IgG2a | Biotin | 1/200 | 135910 | Medical Supply Company, IE |
| Ter-119 | Ter-119 | Rat IgG2b$_{,k}$ | FITC | 1/250 | 557915 | BD, UK |
| CD31 | MEC13.3 | Rat IgG2a | FITC | 1/50 | 102508 | eBioscience, UK |
| CD45 | 30-F11 | Rat IgG2b | FITC | 1/500 | 553080 | BD, UK |
| Lineage | 17A2 | Rat | FITC | 1/10 | 22-7770-72 | eBioscience, UK |
| Sca-1 | D7 | Rat IgG2a$_{,k}$ | FITC | 1/100 | 557405 | BD Biosciences, Oxford, UK |
| gp38 (Podoplanin) | 8.1.1 | Hamster IgG | PE | 1/20 | 12-5381-82 | eBioscience,UK |
| CD73 | TY23 | Rat IgG2a | PE | 1/20 | 550741 | BD, UK |
| PDFGR-α | APA5 | Rat IgG2a | PE | 1.25/100 | 135910 | BD, UK |
| MHC-II | AF6-88.5 | Rat IgG2a | PE | 1/1000 | 553570 | BD, UK |
| CD54 | YN1/1.7.4 | Rat IgG2b$_{,k}$ | PE | 0.625/100 | 12054181 | Medical Supply Company, IE |
| gp38 | 8.1.1 | Mouse IgG | PE | 1/200 | 12-5381-82 | eBioscience,UK |
| CD135 | A2F10 | Rat IgG2a$_{,k}$ | PE | 1/100 | 12-1351 | eBioscience, UK |
| CD16/32 | 2.4G2 | Rat IgG2b,k | PE | 1/200 | 553145 | BD Biosciences, Oxford, UK |
| CD86 | GL1 | Rat IgG2a | PE-Cy7 | 1/20 | 560582 | BD, UK |
| CD31 | 390 | Rat IgG2a$_{,k}$ | PE-Cy7 | 1/50 | 102417 | Medical Supply Company, IE |
| CD48 | HM48-1 | Hamster IgG1$_{,\lambda 3}$ | PE-Cy7 | 1/200 | 560731 | BD Biosciences, Oxford, UK |
| CD41 | MWReg30 | Rat IgG1$_{,k}$ | PE-Cy7 | 1/200 | 133915 | Medical Supply Company, IE |
| c-kit | 2B8 | Rat IgG2b$_{,k}$ | PE-CF594 | 1/400 | 562417 | BD Biosciences, Oxford, UK |
| CD34 | HM34 | Hamster IgG | PerCP-Cy5.5 | 1/50 | 128607 | Medical Supply Company |
| SDC2 | 305515 | Rat IgG2b | APC | 1/50 | FAB2965A | Bio-teche(R&D systems),UK |
| CD44 | IM7 | Rat IgG 2b | APC | 1/100 | 559250 | BD, UK |
| Sca-1 | D7 | Rat IgG2a | APC | 1/100 | 17-5981-81 | eBioscience, UK |
| F4/80 | BM8 | Rat IgG2a | APC | 1/20 | 123116 | Medical Supply Company, IE |
| Sca-1 | D7 | Rat IgG2a | APC | 1/100 | 17-5981-81 | eBioscience, UK |
| c-kit | 2B8 | Rat IgG2b,k | APC | 1/400 | 553356 | BD Biosciences, Oxford, UK |
| CD11b | M1/70 | Rat IgG2b | APC-Cy7 | 1/50 | 557657 | BD, UK |
| CD150 | Q38-480 | Rat IgG2a$_{,k}$ | V421 | 1/50 | 562811 | BD Biosciences, Oxford, UK |
| IL-7R | SB/199 | Rat IgG2b,k | V421 | 1/50 | 562959 | BD Biosciences, Oxford, UK |
| CD45 | 30-F11 | Rat IgG2b | V450 | 1/500 | 560501 | BD, UK |
| Viability Dye | | | eFluor780 | 1/1000 | 65-0865-18 | eBioscience, UK |
| Streptavidin | | | PE | 1/400 | 554061 | BD, UK |
| Streptavidin | | | PerCP | 1/400 | 405213 | Medical Supply Company, IE |
| Streptavidin | | | V421 | 1/400 | 563259 | BD, UK |
| Streptavidin | | | V510 | 1/400 | 405233 | Medical Supply Company, IE |
| Isotype | A95-1 | Rat IgG2b | Purified | 1/100 | 560457 | BD,UK |
| Isotype | | Sheep IgG | Purified | 1/50 | 31243 | Thermofisher, UK |
| Isotype | RTK2758 | Rat IgG2a | Biotin | 1/200 | 400503 | Medical Supply Company |
| Isotype | | Goat IgG1 | Biotin | 1/1000 | A10519 | Thermofisher, UK |
| Isotype | A95-1 | Rat IgG2b | FITC | 1/250, 1/500 | 553988 | BD,UK |
| Isotype | R35-95 | Rat IgG2a$_{,k}$ | FITC | 1/100 | 553929 | BD Biosciences, Oxford, UK |
| Isotype | | Hamster IgG | PE | 1/200 | 12-4914-81 | eBioscience,UK |
| Isotype | A85-1 | Rat IgG1 | PE | 1/20 | 550083 | BD,UK |
| Isotype | A95-1 | Rat IgG2b | PE | 0.625/100 | 553989 | BD,UK |
| Isotype | eBR2a | Rat IgG2a | PE | 1.25/100 | 12-4321-41 | eBioscience,UK |
| Isotype | eBR2a | Rat IgG2a$_{,K}$ | PE | 1/100 | 01124321 | eBioscience, UK |
| Isotype | eB149/10H5 | Rat IgG2b$_{,k}$ | PE | 1/200 | 01124031 | eBioscience, UK |
| Isotype | A95-1 | Rat IgG2b | PE-CF594 | 1/400 | 562308 | BD Biosciences, Oxford, UK |
| Isotype | HTK888 | Hamster IgG | PerCP-Cy5.5 | 1/50 | 400932 | Medical Supply Company |
| Isotype | G235-2356 | Hamster IgG | PerCP-Cy5.5 | 1/50 | 557798 | BD Biosciences, Oxford, UK |
| Isotype | RTK2758 | Rat IgG2a | PE-Cy7 | 1/20 | 400521 | Medical Supply Company |
| Isotype | RTK2071 | Rat IgG1$_{,k}$ | PE-Cy7 | 1/200 | 400415 | Medical Supply Company, IE |
| Isotype | | Hamster IgG1 | PE-Cy7 | 1/200 | 01124914 | eBioscience, UK |
| Isotype | A95-1 | Rat IgG2b | APC | 1/100 | 556924 | BD,UK |
| Isotype | A95-1 | Rat IgG2b | APC | 1/50, 1/400 | 553991 | BD,UK |
| Isotype | eBR2a | Rat IgG2a | APC | 1/100 | 17-4321 | eBioscience,UK |
| Isotype | A95-1 | Rat IgG2b | APC-Cy7 | 1/50 | 552773 | BD,UK |
| Isotype | R35-95 | Rat IgG2a$_{,K}$ | V421 | 1/50 | 562602 | BD Biosciences, Oxford, UK |
| Isotype | R35-38 | Rat IgG2b$_{,k}$ | V421 | 1/50 | 562603 | BD Biosciences, Oxford, UK |
| Isotype | R35-38 | Rat IgG2b | V450 | 1/500 | 562603 | BD,UK |
| Brilliant Stain Buffer | | | | | 563794 | BD, UK |
| CaliBRITE unlabelled beads | | | | | 40486 | BD Biosciences, Oxford, UK |

**Supplemental Table 5. List of Antibodies**

The table contains the reagents used for flow cytometry and histological analysis of HSCs, SCs and MSCs.

| Breeding | % fap+ |
|---|---|
| Breeding 1 | 38.10 |
| Breeding 2 | 47.06 |
| Breeding 3 | 50.00 |
| Breeding 4 | 64.52 |
| Breeding 5 | 52.94 |
| Breeding 6 | 66.67 |
| Breeding 7 | 61.54 |
| Breeding 8 | 36.17 |
| Breeding 9 | 56.00 |
| Breeding 10 | 58.93 |
| Breeding 11 | 41.18 |

52.10 ±10.57 %
42-63%

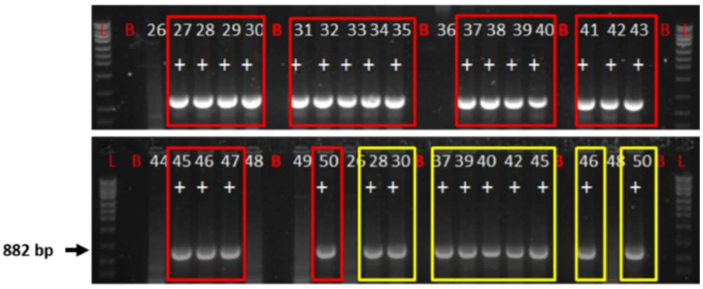


**Supplemental Figure 1**. **Representative *fap* genotype by digesting ear biopsies.**
The table contains the average of *fap* expression over the breeding. Representative agarose gel of the genotyping process for 25 mice detected by PCR. The modified gene fraction sizes at 882 bp (arrow). FAP transgenic mice highlighted with red squares, and the duplicates result in yellow.

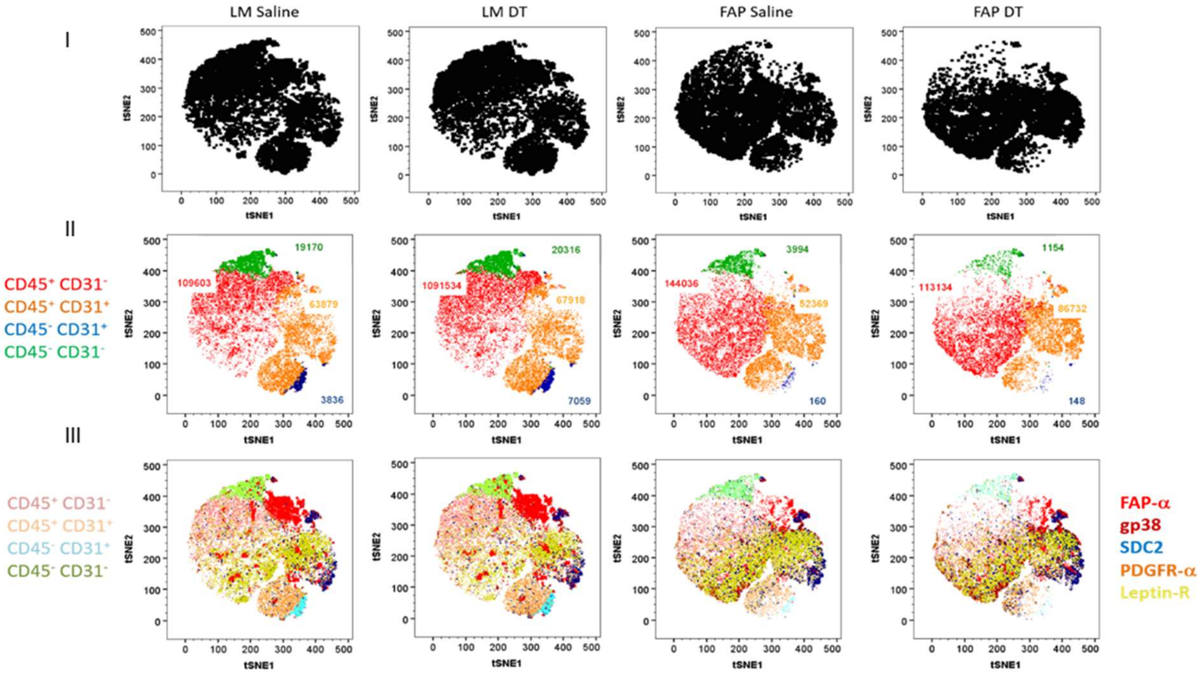


**Supplemental Figure 2**. **Cellular population complexity of BM-FAP$^{DM2}$.**
t-Distributed stochastic neighbour embedding (t-SNE) visualisation of 200,000 events from a representative sample of the study from LM and FAP transgenic mice D3 after DTX or saline administration. **(I)** Distribution map of the alive total cell counts. **(II)** Distribution map of the cellular compartments based on CD45 and CD31 expression: Haematological (red), double-positive (orange), endothelial (red), and stromal clusters (red). **(III)** Distribution map of the cellular subpopulations expressing stromal cell markers FAP, gp38, SDC2, PDGFR-α and Leptin-R within each compartment (light coloured).

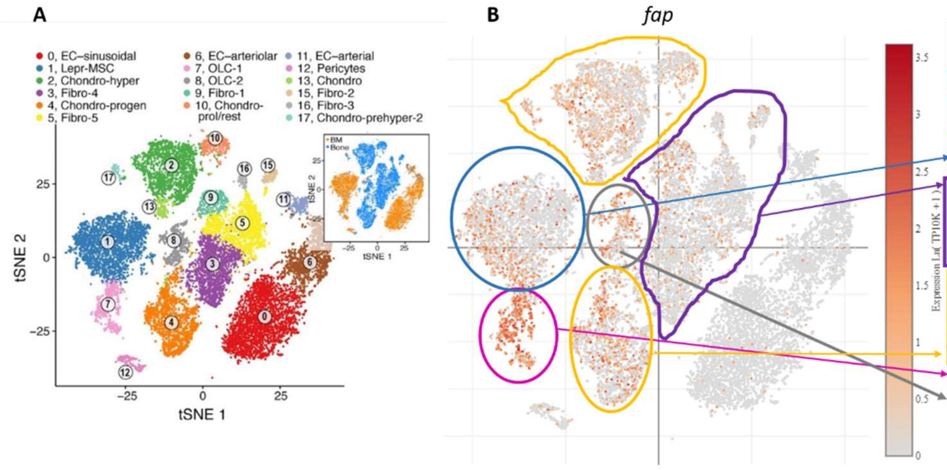


| Cluster name | Cluster number | Total events number | % |
|---|---|---|---|
| EC-sinusoidal | 0 | 3456 | 16.79 |
| EC-arteriolar | 6 | 1793 | 8.71 |
| EC-arterial | 11 | 454 | 2.21 |
| Perycites | 12 | 290 | 1.41 |
| Leptin-R | 1 | 1989 | 9.66 |
| Fibro-1 | 9 | 512 | 2.49 |
| Fibro2 | 15 | 190 | 0.92 |
| Fibro3 | 16 | 163 | 0.79 |
| Fibro4 | 3 | 2090 | 10.15 |
| Fibro-5 | 5 | 1806 | 8.78 |
| Chondro hyper | 2 | 2250 | 10.93 |
| Chondro-proge | 4 | 2025 | 9.84 |
| Chondro-prol/rest | 10 | 508 | 2.47 |
| Chondro | 13 | 277 | 1.35 |
| Chondro-prehyper | 17 | 130 | 0.63 |
| OLC-1 | 7 | 825 | 4.01 |
| OLC-2 | 8 | 722 | 3.51 |

**Supplemental Figure 3. Molecular expression of *fap* in mouse bone marrow stroma**.
**(A)** Seventeen BM stroma cell clusters were taken from Baryawno 2019 study. t-SNE of 20,896 non-hematopoietic cells, annotated post hoc and coloured by clustering, bone or bone marrow location (inset). **(B)** Cell-type marker distribution for *fap* at single-cell RNA expression. Coloured gates and arrows identify clusters where *fap* is expressed. **(C)** The coloured table contains the absolute number of events and the frequency of the seventeen clusters of non-hematopoietic cells in homeostasis.

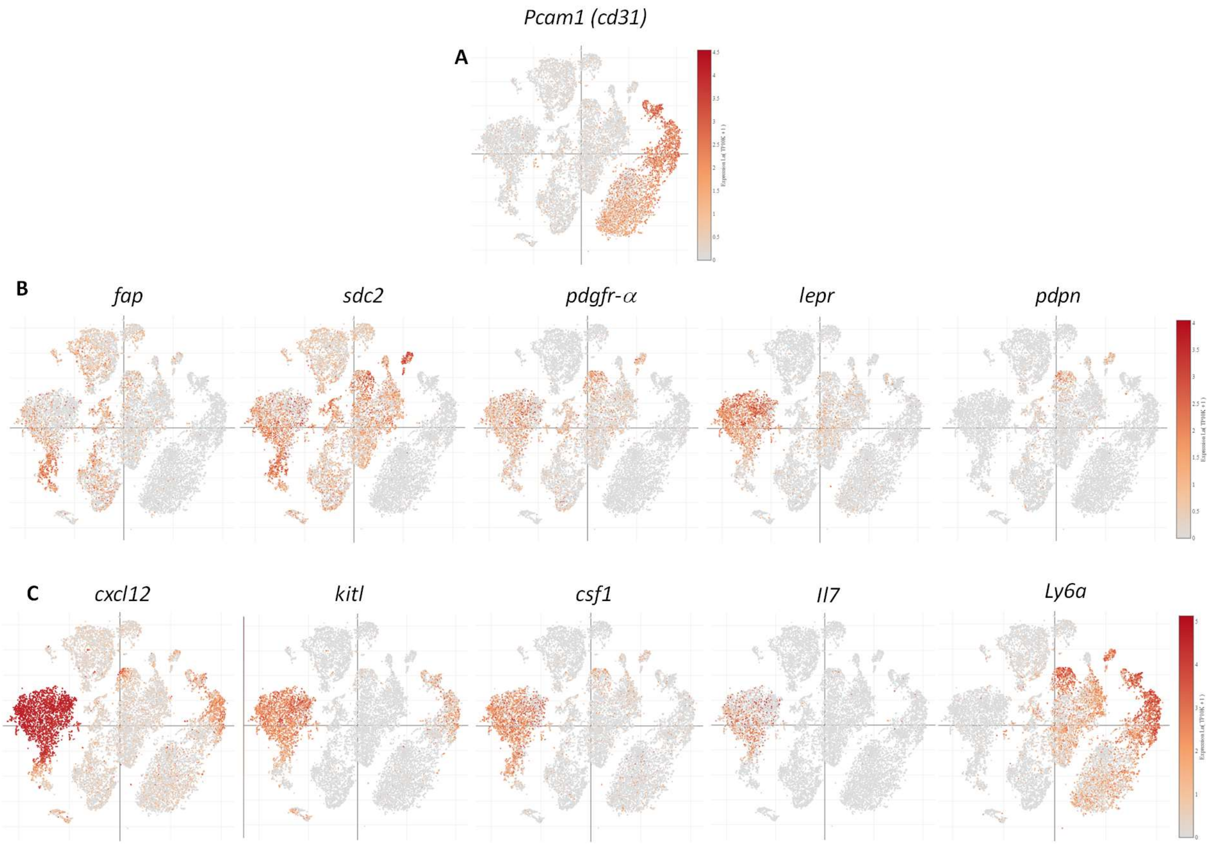


**Supplemental Figure 4. Molecular expression of stromal cell markers and factors in mouse bone marrow stroma.**

Genes corresponding to the cell surface markers analysed in our studies extracted from single-cell RNA expression of mouse bone marrow (Baryawno 2019) contained in the Broad Institute portal. (A) t-SNE of 20,896 non-hematopoietic cells used to localise the expression of the endothelial, (B) stromal cell markers and (C) lineage-specific growth factors.

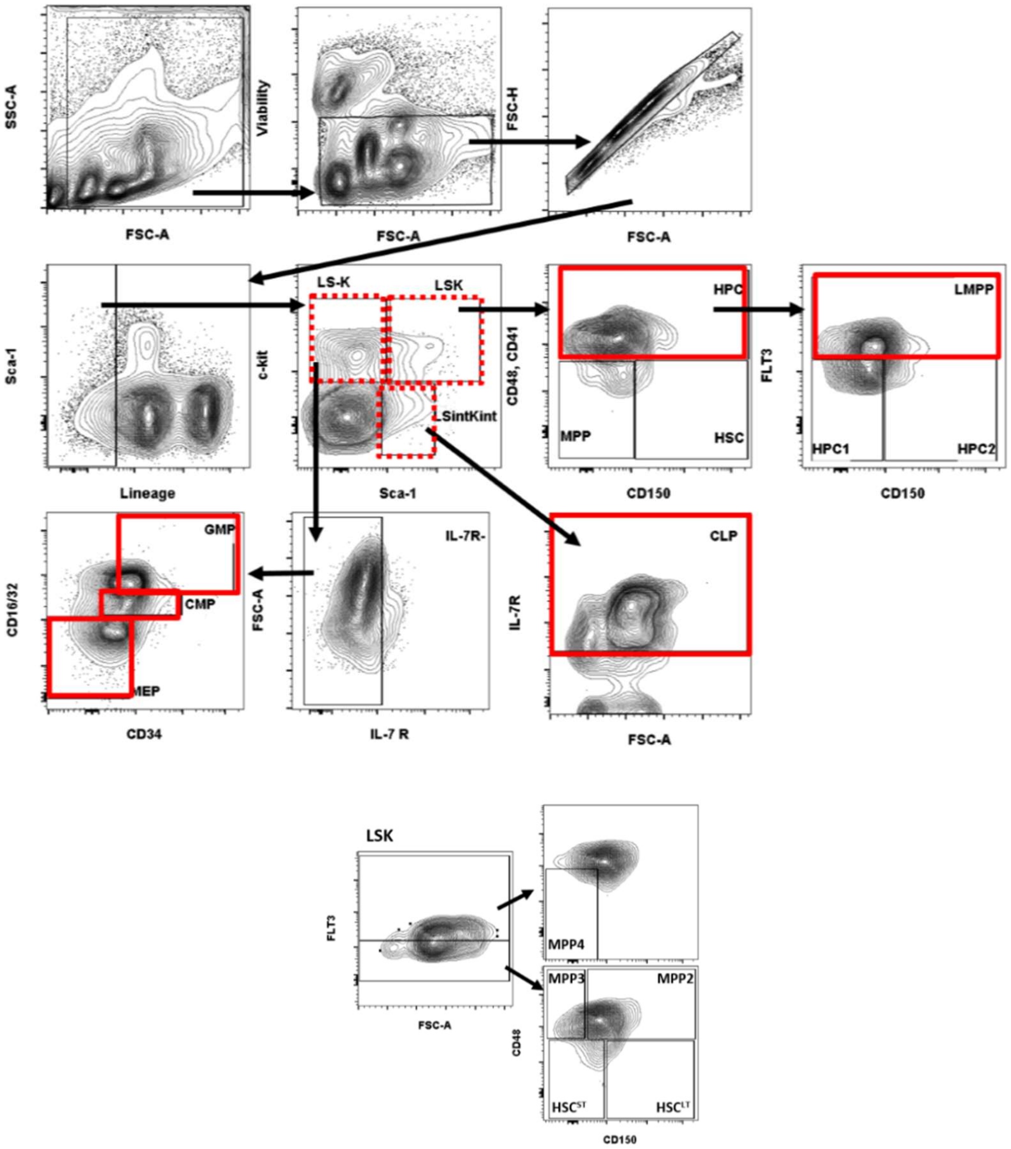


**Supplemental Figure 5. Flow cytometry gate strategy to identify the haematopoietic cell populations.**

Red dotted line used to highlight the intermediated cell populations whilst the red line used to identify the oligopotent progenitors. MPPs subsets and HSC long and short term (HSC$^{LT}$ & HSC$^{ST}$) were gated from LSK cell population.

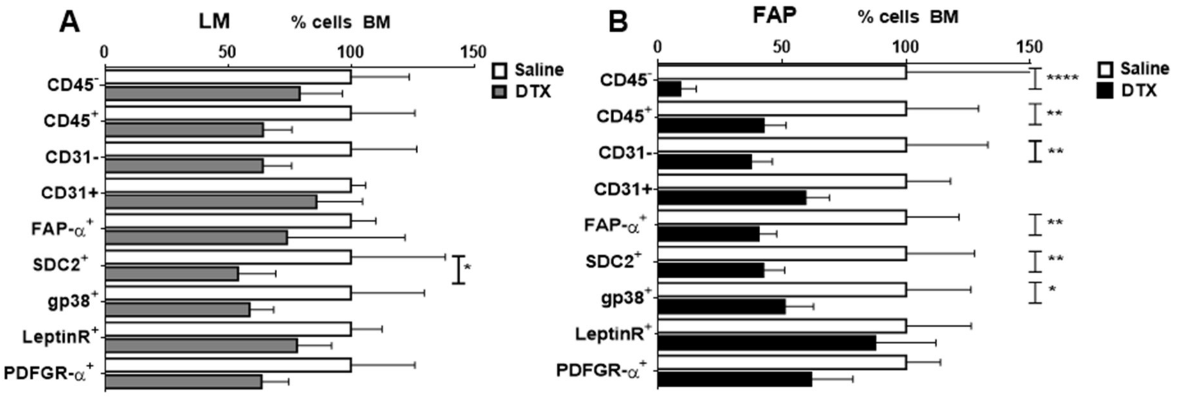


**Supplemental Figure 6**. **Cell surface single proteins expression in BM after FAP-DTX ablation.**

Frequency of single protein in the surface of the BM cells from LM (A) and (B) FAP at D3 after DTX administration or saline as a control. n=4, mean ± SEM, (α=0.05), *,P<0.05; **,P<0.01; ****,P<0.0001).

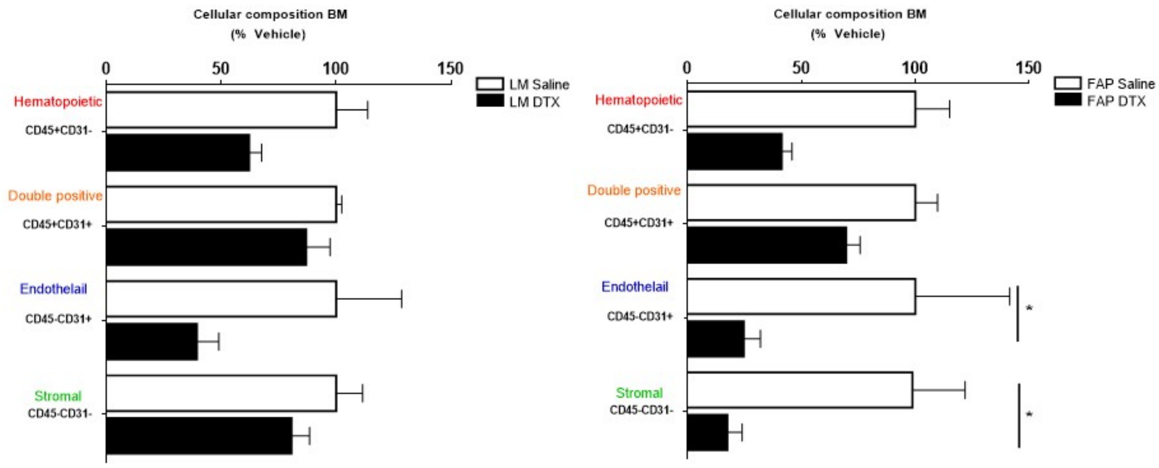


**Supplemental Figure 7. Cellular compartments frequency in BM after FAP-DTX ablation.**

Frequency of cells based on the CD45 and CD31 expression in LM and FAP at D3 after DTX administration. n=4 mean ± SEM, α=0.05*,P<0.05; **,P<0.01.

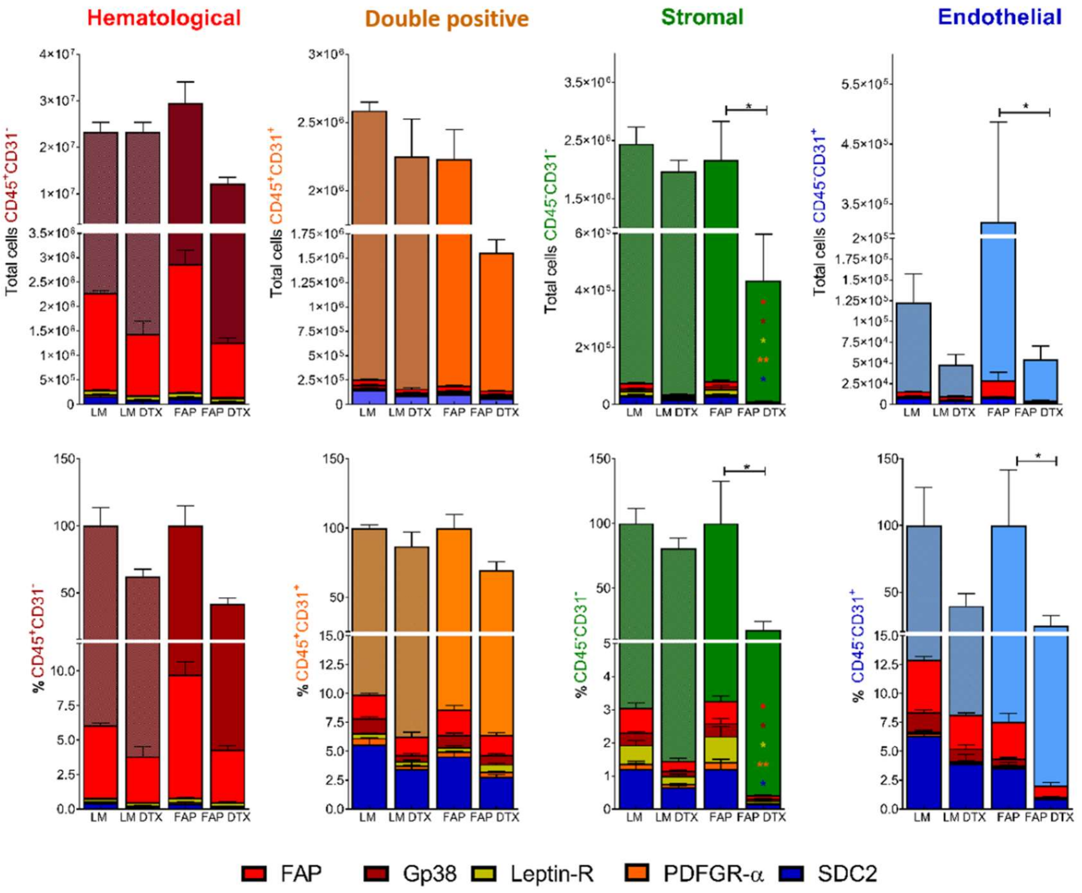


**Supplemental Figure 8**. **Comprehensive census of the cell subpopulations in BM after FAP-DTX ablation.**

The top section contains the grouped-bar graphs with the absolute cell counts per subset and the bottom graphs the calculated frequency for both groups, LM and FAP at D3 after DTX administration or saline as a control. n=4 per group, mean ± SEM; *,$P<0.05$, **,$P<0.01$.

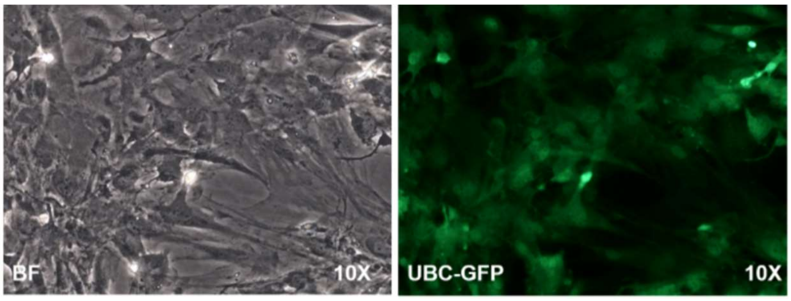


**Supplemental Figure 9. Cell morphology of the expanded BM-UBC-GFP-MSCs**

Representative microscopic visualisation of cell culture and GFP expression of the MSCs at P4. Bright-field and fluorescence image for GFP.

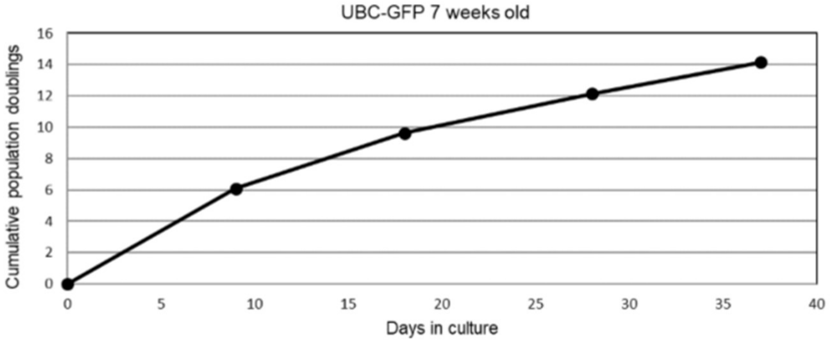


**Supplemental Figure 10. Cell growth curve of the BM-UBC-GFP-MSCs**
The graph contains the mean of the accumulated doublings versus time of BM-UBC-GFP-MSCs batches generated.

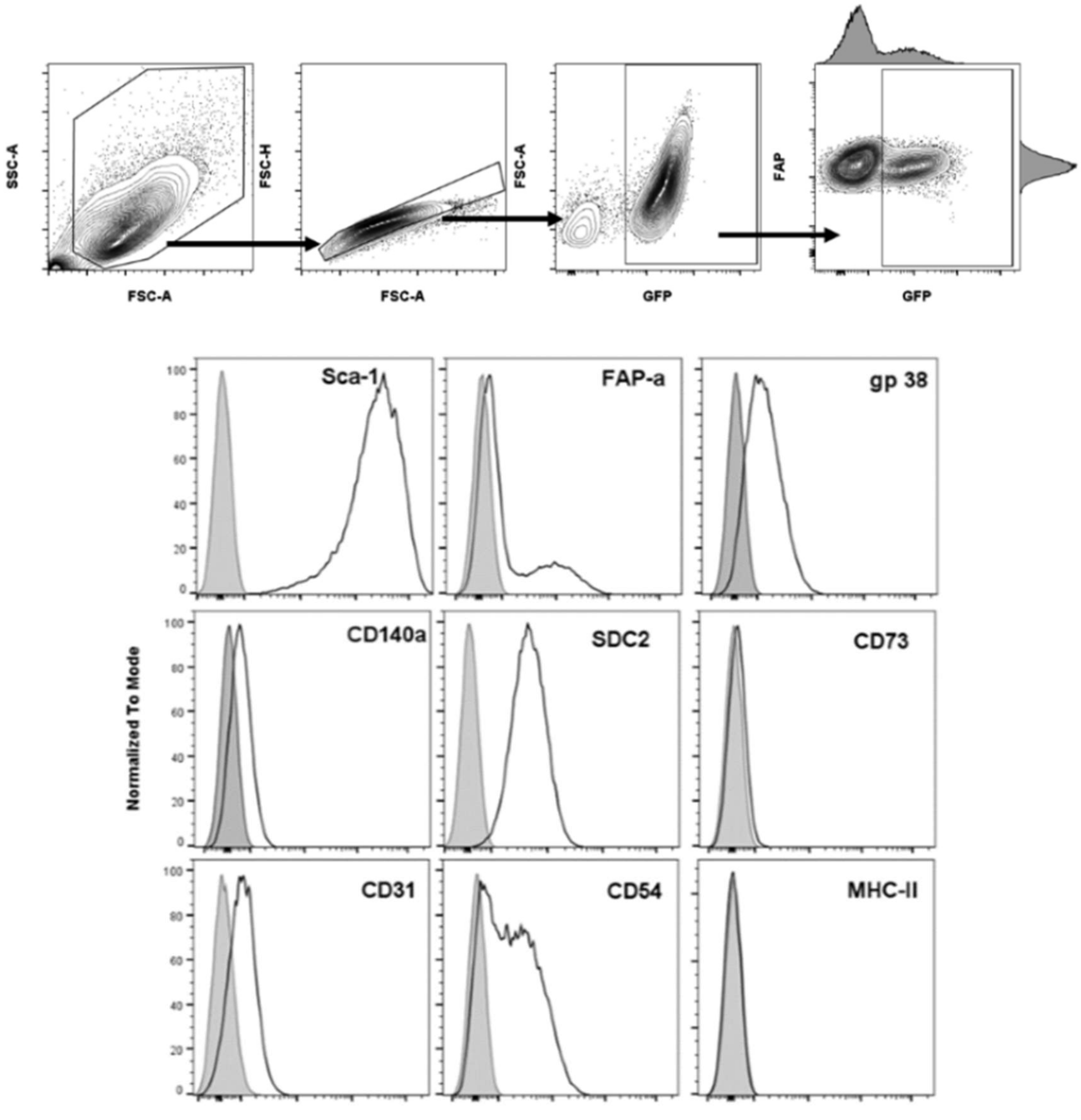


**Supplemental Figure 11. Flow cytometry characterisation of the BM-UBC-GFP-MSCs.**
Gate strategy followed to confirm GFP and FAP expression at P4 before administration. A short panel of cell surface markers was checked to confirm cell viability.
